# Long-term small effective population size, inbreeding, and a recessive lethal haplotype drive premature death in the endangered Devils Hole pupfish (*Cyprinodon diabolis*)

**DOI:** 10.64898/2026.06.06.730634

**Authors:** David Tian, Nicolas Alexandre, Naomi Siu, Vanessa Muhl, Sean C. Lema, Bruce J. Turner, Olin Feuerbacher, Kevin Wilson, Michael Schwemm, Priya Moorjani, Jennifer Gumm, Christopher H. Martin

## Abstract

As anthropogenic habitat fragmentation and population decline accelerate globally, growing numbers of species face compounding demographic and genetic threats to long-term survival. Many populations are already forced to persist at chronically small sizes, yet the genomic and fitness consequences of this fate remain poorly understood. Here we leverage the demographic history of the Devils Hole pupfish to investigate how long-term small population size and recent bottlenecks have shaped genetic diversity, genetic load, inbreeding, and fitness through comparative population genomics, historical sequencing, and sampling embryos that died prematurely during development. We find that genetic diversity in Devils Hole pupfish is among the lowest recorded in the wild and that fixed load is high, consistent with thousands of generations of isolation at small population size. Even in the face of this low diversity and high fixed load, we show that inbreeding is still strongly associated with premature embryonic death, which affects up to 25% of offspring in the captive refuge and can be identified in advance based on a characteristic elongated heart tube and reduced heart rate. We discovered a recessive lethal haplotype segregating at ∼20% frequency that accounts for 50% of embryonic deaths and contains mutations in MIB1 and MMP16, genes associated with cardiomyopathy and atrial fibrillation. Our findings link genotype, phenotype, and fitness in an iconic endangered species to provide a rare comprehensive view into the evolutionary dynamics and consequences of long-term small effective population size, demonstrating that endangered species remain vulnerable to inbreeding depression despite extremely low genetic diversity.

## Introduction

A major goal in conservation biology is to understand the consequences of small population size and the factors and processes that may lead to extinction (1). Small populations are vulnerable to demographic and environmental stochasticity (2) and a range of genetic threats, which include reduced adaptive capacity (3–5), increased inbreeding depression (6–9), and elevated realized load, the portion of the genetic load whose fitness effects are expressed (10–13). Despite these vulnerabilities, some species nevertheless persist at small population size, defying expectations of mutational meltdown, a feedback loop where the accumulation of deleterious alleles causes reductions in fitness and population size, eventually leading to extinction (14–16).

One explanation for this paradox is the purging of strongly deleterious alleles in small populations (17, 18) and the loss of inbreeding load due to drift, which may reduce the extent of inbreeding depression (19–21). However, inbreeding depression may persist despite purging (22) and deleterious alleles can reach high frequency or fixation due to drift in long-term small populations, reducing population mean fitness (23, 24). As a result, the fitness consequences and extinction risks associated with population decline and deleterious variation in wild populations remain dynamic and difficult to predict (25, 26). Identifying and quantifying the genomic and fitness consequences of long-term small effective population sizes is therefore crucial to manage the conservation of endangered species.

The Devils Hole pupfish (*Cyprinodon diabolis*) offers an excellent opportunity to connect the genomic and fitness consequences associated with long-term isolation and small population size. This species is restricted to Devils Hole, a small water-filled fault cavern in a disjunct unit of Death Valley National Park located in Amargosa Valley, NV that is the smallest known habitat for a vertebrate in the world (27). Devils Hole pupfish, along with *C. nevadensis* and *C. salinus* pupfishes endemic to Death Valley National Park and Ash Meadows Wildlife Refuge, are estimated to have become isolated at the end of the Pleistocene 10-20 kya, when a large network of pluvial lakes and interconnecting streams in Death Valley began to dry (28–30).

The Devils Hole pupfish population declined in the early 1970s due to nearby groundwater extraction that lowered water levels within Devils Hole, before the U.S. Supreme Court intervened and mandated a minimum water level (31). Afterwards, the population recovered and remained stable at around 200-500 individuals in the spring and fall, respectively (32). However, the population began to decline again in the mid-1990s, reaching all-time lows of 38, 35, and 38 in 2007, 2013, and 2025, respectively. Following the 2013 bottleneck, a captive refuge at Ash Meadows Fish Conservation Facility was established to safeguard against extinction of the wild population. The captive population fluctuates and is supplemented with eggs collected monthly from the wild that are reared in captivity, reaching over 500 individuals in 2023. However, a proportion of developing embryos in the captive population, up to one quarter in lab-based breeding groups, do not survive to adulthood.

Here, we investigate the genomic and fitness consequences of long-term isolation and small population size in the Devils Hole pupfish. We assembled the first chromosome-scale *de novo* genome assemblies for the Devils Hole pupfish and Ash Meadows pupfish, along with population-level whole-genome resequencing of 186 Death Valley pupfishes and outgroups, including 78 wild, captive, historical, and embryonic lethal Devils Hole pupfish samples. We examine how demographic history has shaped population structure, genetic diversity, and genetic load in Devils Hole pupfish relative to neighboring Death Valley pupfish species. Then, we estimate the fitness consequences of inbreeding and identify a recessive lethal haplotype associated with embryonic lethality and a known heart defect phenotype in Devils Hole pupfish embryos. Our results highlight how small populations with reduced inbreeding load can still be vulnerable to inbreeding depression due to elevated frequencies of recessive lethal alleles.

## Results

### Genome assembly

Using PacBio HiFi data and Dovetail Omni-C data, we generated *de novo* genome assemblies for *C. diabolis* (UCB_CyDiab_1.0) and *C. nevadensis mionectes* (UCB_CyNevMio_1.0). Both assemblies were 1.22 Gb in length and highly contiguous with similar numbers of scaffolds (*diabolis*: 1102, *mionectes*: 1726), scaffold N50 (*diabolis*: 48.0 Mb, *mionectes*: 46.3 Mb), and scaffold L75 (*diabolis*: 19, *mionectes:* 20) (Supp. Table 1). In addition, both assemblies were high quality with overall base pair quality values of 61. Both assemblies had a GC content of 39.5% with similar proportions of repetitive regions, with 44.1% and 46.1% of the *diabolis* and *mionectes* reference genomes soft-masked, respectively. Both assemblies showed high k-mer completeness values (*diabolis*: 99%, *mionectes*: 94%) and nearly all *Actinopterygii* complete and single-copy gene orthologs were present in the assemblies (*diabolis*: 95%, *mionectes*: 95%) based on BUSCO (33) (Supp. Table 2). We used the high-quality *Fundulus heteroclitis*genome (GCA_011125445.2) to name our chromosomes.

### Population structure

Principal component and admixture analyses based on 683,969 LD-pruned SNPs from a total of 11,199,716 high-quality SNPs indicated the presence of strong population structure among Death Valley pupfishes and outgroups, consistent with previous results (30, 34) (Fig. 1B and 1C). The small amount of *C. diabolis* ancestry within *C. nevadensis mionectes* may be explained by a documented translocation of 75 Devils Hole pupfish into an unspecified nearby spring in Ash Meadows in 1946 (35, 36). Several *C. diabolis* historical specimens or poor-quality samples from deceased individuals found in Devils Hole showed putative signatures of admixture, but this is likely due to low coverage, DNA degradation, or contamination; these individuals were removed from subsequent analyses (Fig. 1C).

**Fig. 1.**
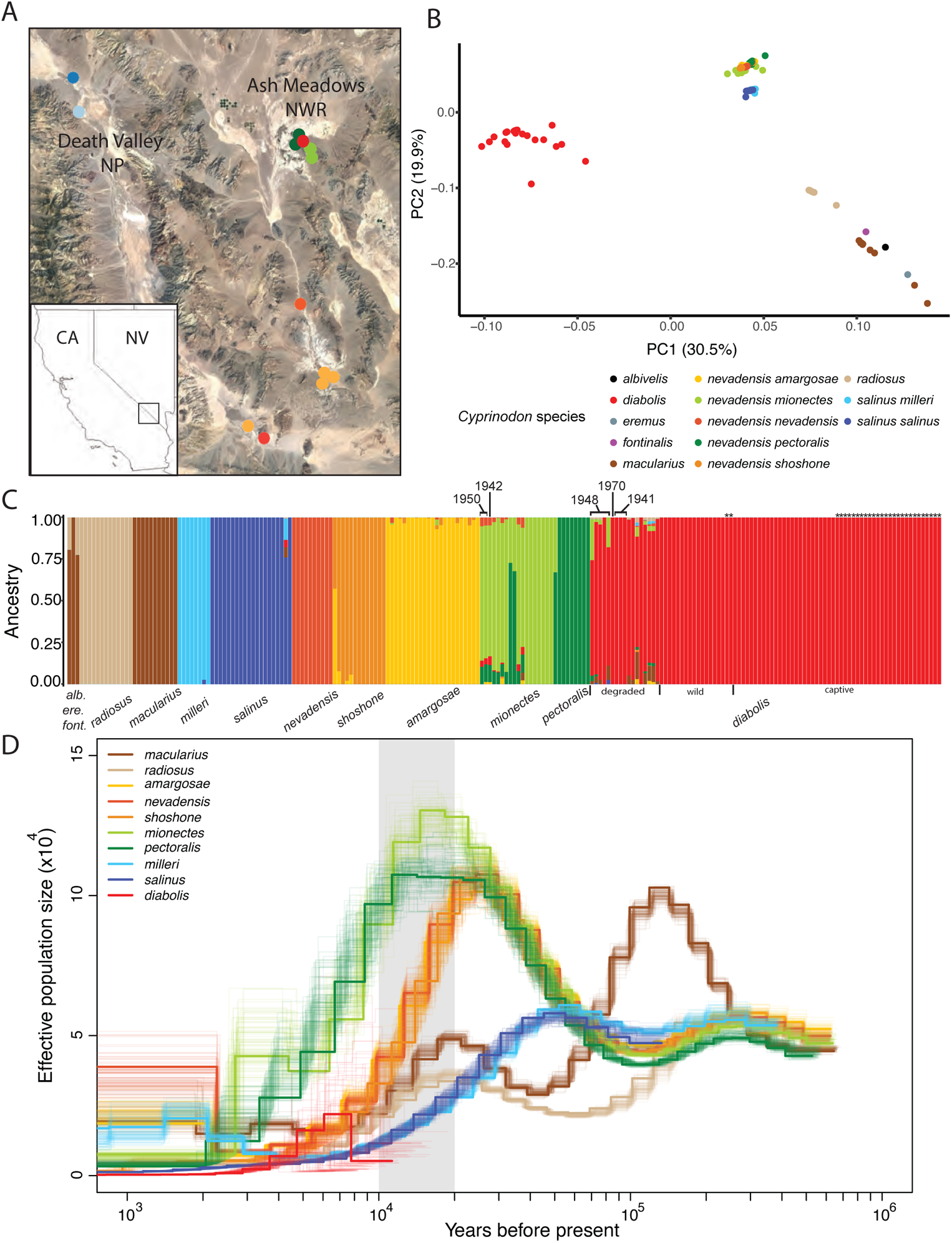
**Population structure and historical demography of desert pupfishes**. (A) Map of pupfish species sampling locations in Death Valley and Ash Meadows National Wildlife Refuge. Inset map shows location of Death Valley along the CA and NV border. Outgroups not shown. (B) Principle component analysis of desert pupfishes indicating considerable population structure among species. (C) Ancestry proportions for individuals in Death Valley National Park, Ash Meadows National Wildlife Refuge, and outgroup desert pupfishes (*C. albivelis, C. eremus, C. fontinalis, C. macularius, C. radiosus*) estimated from a LD-pruned SNP dataset in ADMIXTURE (k = 10). Colors in both PCA and ADMIXTURE represent individuals from different species or populations. Principle component and admixture analyses based on 683,969 LD-pruned SNPs out of a total 11,199,716 SNPs. Stars above ancestry columns of Devils Hole pupfish individuals mark embryonic lethal samples. (D) PSMC trajectories and 100 bootstraps showing historical effective population sizes inferred from a representative high-coverage individual per species. To scale the axes, we used a mutation rate of 8.09 x 10^-9^ per bp per generation and a generation time of 0.75 years. The grey bar marks when pluvial lakes in Death Valley began to dry up 10 - 20 kya. The lack of resolution for *C. diabolis* beyond a few thousand years may be due to insufficient older coalescent events under long-term small population size and extreme bottlenecks.

### Historical demography and effective population size

To better understand the demographic history of Death Valley pupfishes, we inferred an admixture graph and long-term effective population size trajectories using the pairwise sequentially Markovian coalescent (PSMC) model across all desert pupfish populations (37) (Fig. 1). We found evidence of extensive historical admixture across desert pupfishes, consistent with a historically interconnected aquatic landscape (Supp. Fig. 1), but strong population structure today (Fig. 1 B, C). Effective population size trajectories indicate that *C. nevadensis* and *C. salinus* populations diverged ∼50,000 years ago as *C. salinus* populations began to decline and that *C. nevadensis* populations began to decline 10 – 20 kya, coinciding with the drying of the Mojave Desert and Lake Manly in Death Valley towards the end of the Pleistocene (29) (Fig. 1D).

For *C. diabolis*, effective population size was consistently low over the course of the last few thousand years. However, the PSMC model was unable to infer effective population sizes further back in time due to lack of power, potentially due to the initial colonization of Devils Hole or past bottlenecks that obscure deeper coalescent events (Fig. 1D). This pattern was supported by increasingly stochastic bootstraps further back in time and was consistent across multiple individuals (Supp. Fig. 2,3). We applied PSMC to two high coverage *C. diabolis* historical genomes that suggest the population may have become isolated around 10,000 years ago (Supp. Fig. 4). Given these limitations, we consequently also estimated long-term effective population size defined here as *N_e_* = π/(4μ) to be 426 for Devils Hole pupfish, based on a heterozygosity of 1.38 x 10^-5^ (Fig. 2B) and inferred mutation rate of 8.09 x 10^-9^ bp^-1^ gen^-1^ estimated for the Devils Hole pupfish (38). We estimated contemporary effective population size (2020-2022) to be 18 (90% CI: 13.67 – 23.70) based on patterns of linkage-disequilibrium (39) (Supp. Table 3).

**Fig. 2.**
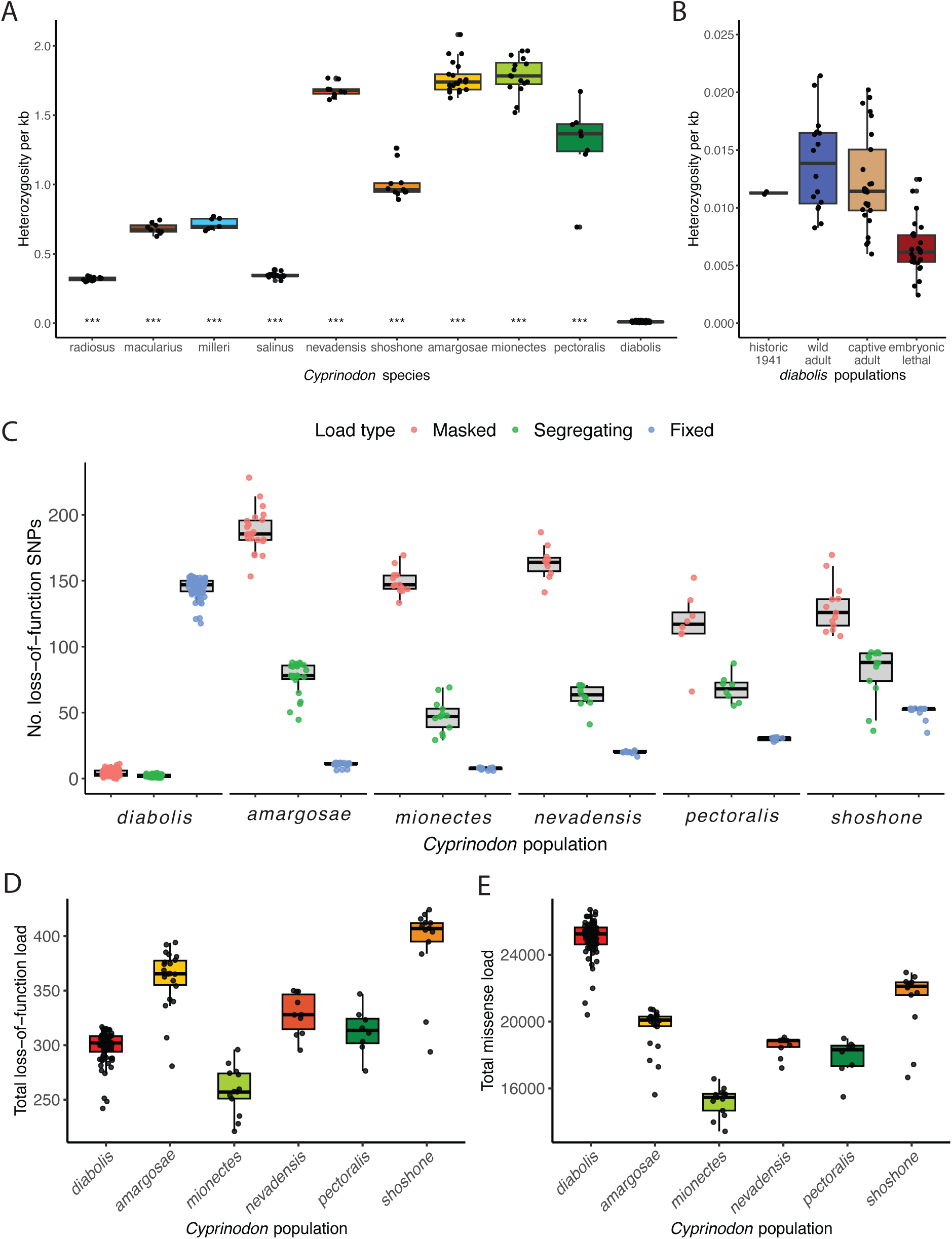
**Heterozygosity and genetic load across populations. (**A) Heterozygosity per species with stars representing significant differences (*P* < 0.001) in heterozygosity for each species relative to *C. diabolis* (Mann-Whitney tests). (B) Comparison of heterozygosity across historic, wild, captive, and embryonic lethal populations of *C. diabolis.* (C) Individual loss-of-function mutation loads based on number of putatively deleterious alleles detected in each individual for Devils Hole pupfish and five *C. nevadensis* populations. These mutations were categorized into masked (heterozygous SNPs) and realized load (homozygous SNPs). Realized load was further parsed into segregating (variable within a species) and drift load (fixed within a species or population). Differences in load between species are relative rather than absolute, given that we were unable to comprehensively polarize our VCF. (D) Total loss-of-function load was calculated as the number of derived loss-of-function alleles for each individual. (E) Total missense load was calculated as the number of derived missense alleles for each individual.

**Fig. 3.**
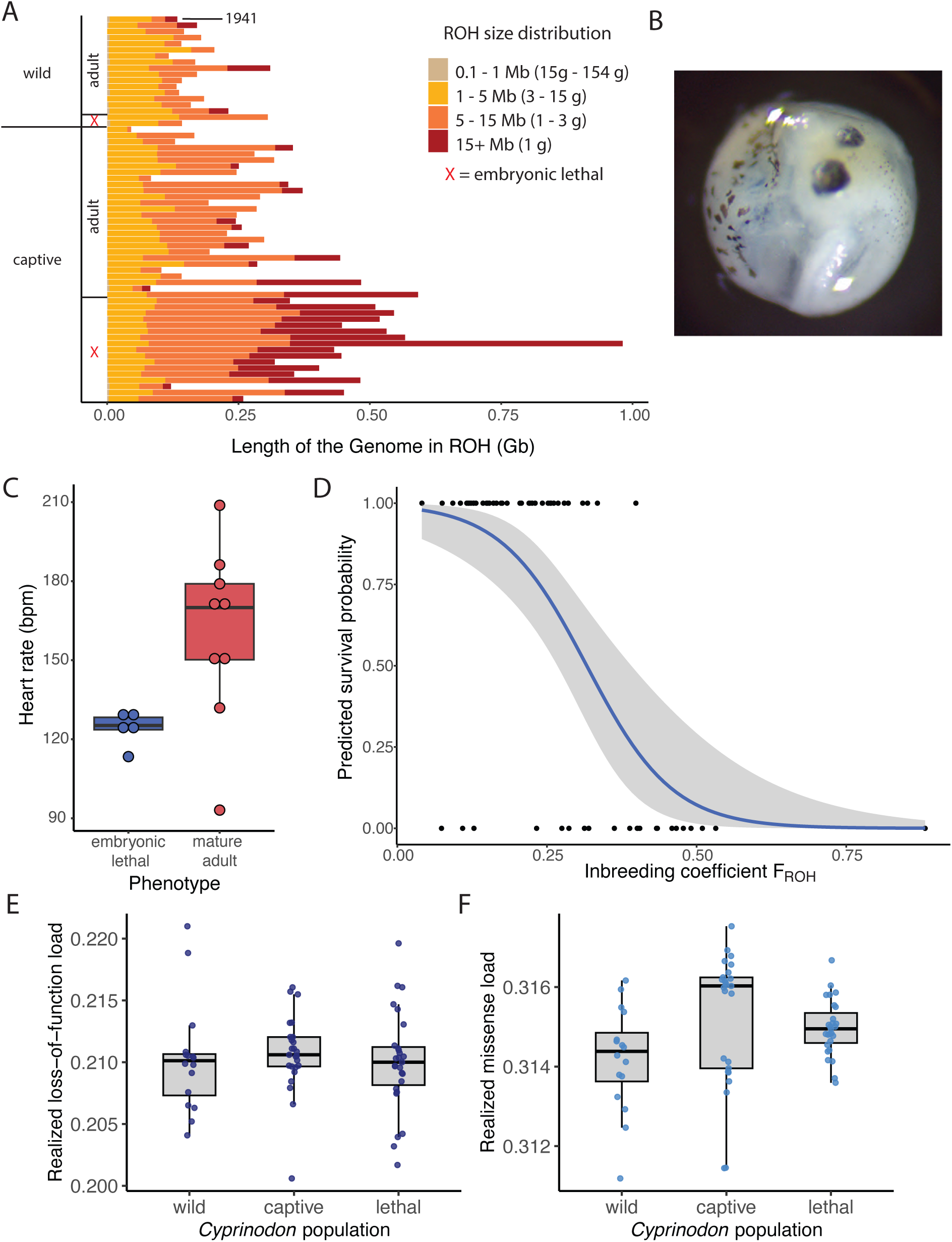
Inbreeding depression and reduced heart rate in embryonic lethal Devils Hole pupfish. (A) Distribution of short (0.1 Mb – 1 Mb) – very long (15+ Mb) runs of homozygosity in Devils Hole pupfish. Each row represents an individual with red Xs designating embryos that died prematurely during development (embryonic lethal). Note that we included one historical genome from 1941. (B) Dorsal view of an embryonic lethal Devils Hole pupfish. (C) Heart rate (beats per minute) at 5 days-post-fertilization in embryos that died prematurely or survived to adulthood. (D) Generalized linear mixed model of survival probability and 95% confidence interval over a range of inbreeding coefficients (*F_ROH_*). (E) Realized loss-of-function load per individual normalized by number of called genotypes across wild adult, captive adult, and embryonic lethal Devils Hole pupfish. (F) Realized missense load per individual normalized by number of called genotypes across wild adult, captive adult, and embryonic lethal Devils Hole pupfish.

**Figure 4.**
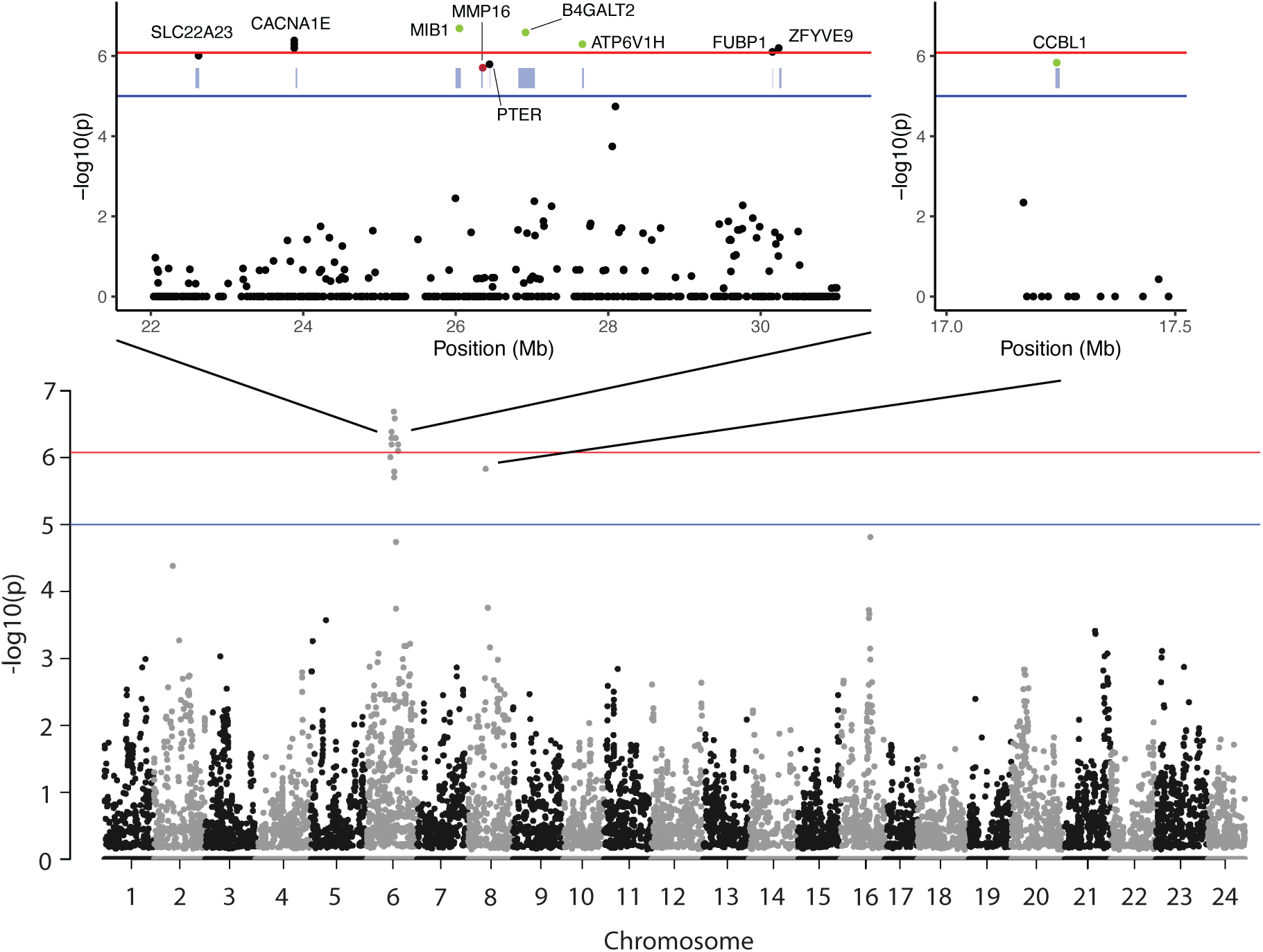
Identification of a candidate recessive lethal haplotype driving embryonic lethality. Genome-wide association analysis testing for recessive lethal alleles associated with embryonic lethality, with blue line representing genome-wide significance threshold of *P* < 1 x 10^-5^ and red line representing Bonferroni-corrected threshold of *P* < 8.29 x 10^-7^. Insets show outlier SNPs on chromosomes 6 and 8. Purple rectangle represent the closest gene to each significant SNP. Among significant SNPs: black, intergenic; green, intronic; red, exonic.

### Genetic diversity

We found low heterozygosity across all desert pupfish populations, consistent with their small population sizes and isolation following the drying of the Mojave Desert and corroborating previous mitochondrial (40–42), retrotransposon (43), and RAD-seq analyses (34) (Fig. 2A). *C. nevadensis* populations all showed similar relative levels of heterozygosity, although *C. nev. shoshone* had lower genetic diversity, consistent with a reintroduction of 70 captive-bred individuals all descended from a single female into the spring a few years prior to sampling (BJT pers. obs.). *C. macularius* and *C. salinus* populations had lower heterozygosity, consistent with their lower historical effective population size (Fig. 1D). *C. radiosus* also showed very low heterozygosity, consistent with the fact that all existing individuals today are descended from a single population of roughly 400 individuals that were saved from a drying spring in 1969 by P. Pister (44).

However, *C. diabolis* had the lowest wild heterozygosity (mean *H* = 1.38 x 10^-5^) of all populations, with on average only one heterozygous site per 77 kb, on par with the San Nicolas Island population of Channel Island foxes, one of the lowest estimates of genetic diversity in the wild to date (20, 45, 46). Across the genome, heterozygosity was uniformly low, consistent with long-term small population size (Supp. Fig. 5). Across Devils Hole pupfish populations, the captive refuge population contained less genetic diversity than the wild population (2019-2020) despite regular introductions of individuals derived from wild-collected eggs and a similar census population size, although this difference was not significant (*P* = 0.49, Tukey’s HSD test). Embryonic lethal individuals had significantly lower heterozygosity than wild and captive adults (wild vs. embryonic lethal *P* = 4.30 x 10^-8^, captive vs. embryonic lethal *P* = 8.26 x 10^-7^, Tukey’s HSD test). Finally, we found no significant difference in heterozygosity between our historic (1941) samples and modern (2019-2020) wild samples, although we were limited to only two higher coverage historic samples (Fig. 2B).

### Genetic load

We leveraged our *C. salinus* and outgroup (*radiosus, macularius, albivelis, eremus, fontinalis*) samples to polarize alleles and decompose genetic load into loss-of-function vs. missense load and masked vs. realized load across Devils Hole pupfish and the other five *C. nevadensis* populations (Fig. 2 C, D, E). We were unable to confidently polarize all variants; thus, the genetic load results we present represent relative rather than absolute differences between populations. Devils Hole pupfish had intermediate total loss-of-function load (298.4 ± 15.1 SD loss-of-function deleterious alleles per individual) relative to other *C. nevadensis* populations but significantly higher missense load (24,997.9 ± 1118.1 SD missense deleterious alleles per individual) relative to other *C. nevadensis* populations (*diabolis* x other *nevadensis* populations, all *P* < 0.0001, Tukey’s HSD) (Fig. 2 D,E).

The composition of genetic load contrasted starkly between Devils Hole pupfish and other *C. nevadensis* populations. The proportion of loss-of-function load that was masked (inbreeding load) was substantially lower in Devils Hole pupfish (mean = 1.39%) compared to *C. nevadensis* populations (range = *shoshone:* 33.67% - *mionectes*: 58.05%) (Fig. 2C). Conversely, Devils Hole pupfish had substantially higher proportions of realized loss-of-function load (mean = 98.61%) compared to *C. nevadensis* populations. We further decomposed realized load into homozygous deleterious alleles fixed within the population (drift load) versus not fixed (segregating load), and found that the vast majority of realized load in Devils Hole pupfish was fixed (mean = 98.54%) whereas *C. nevadensis* populations had much lower proportions of realized load fixed within the population (range = *amargosae:* 12.51% - *shoshone*: 39.31%).

Among the 154 loss-of-function variants fixed in Devils Hole pupfish in our high-quality modern samples, we searched our two high quality historical genomes for the same genotypes. We only found evidence of one modern fixed variant that was segregating in historical samples (1948), suggesting that nearly all drift load accumulated over longer timescales due to small population size rather than the recent period of intensive human management and extreme population bottlenecks. Similar patterns of low masked load and high realized load in Devils Hole pupfish relative to other *C. nevadensis* populations were observed for missense alleles (Supp. Fig. 6).

Finally, we identified a total of 37 fixed, loss-of-function variants unique to Devils Hole pupfish and found in no other samples in our dataset (Supp. Table. 4). Among these variants, we found no significant enrichment of any gene ontology (KEGG) categories, consistent with the expectation of random deleterious alleles drifting to fixation. We also identified two segregating deletions unique to Devil Hole pupfish: PFKM, a kinase that serves as a regulatory enzyme of glycolysis (47, 48) and RAD23A, a protein involved in nucleotide excision repair and protein degradation (49–51) (Supp. Table. 5).

### Inbreeding and inbreeding depression

Consistent with our heterozygosity results, captive adults were on average more inbred than wild adults but not significantly so (wild adult F_ROH_ = 0.16 vs. captive adult F_ROH_ = 0.22; *P* = 0.33, Tukey’s HSD). In addition, there was an apparent increase in inbreeding over time in the wild population (mean wild adult F_ROH_ = 0.16, 95% CI: 0.13 – 0.18) relative to a single high-coverage historical museum specimen from 1941 that had a F_ROH_ of 0.12.

### Premature embryonic death during development

Depending on the collection period and source, between 7-25% of developing embryos collected in captivity and tracked in individual culture dishes died at 5 days post-fertilization (dpf), with some embryos displaying a characteristic “stringy heart” phenotype. This observation in conjunction with our finding of reduced heterozygosity in embryonic lethal individuals relative to mature adults (Fig. 2B) led us to hypothesize that Devils Hole pupfish may be suffering from inbreeding depression. Consequently, we measured heart rates among developing embryos from videos and inbreeding among Devils Hole pupfish from the proportion of the genome made up of runs of homozygosity (ROH) that were at least 100 kb long (Fig. 3).

On average, embryos with the heart defect phenotype had a 23% slower heart rate than healthy embryos that developed into adults (Fig. 3C; 124.2 bpm vs. 160.3 bpm; *P* = 0.013, two-sample *t*-test). In addition, captive embryonic lethal individuals were significantly more inbred than captive adults (captive adult F_ROH_ = 0.22 vs. captive embryonic lethal F_ROH_ = 0.40; *P* = 1 x 10^-5^, Tukey’s HSD) (Fig. 3A) and had significantly more long ROHs (> 15 Mb) than mature adults (two-sample t-test, *P* = 7.98 x 10^-8^), indicative of recent inbreeding within the past generation. Overall, a 10% increase in inbreeding resulted in a 75% decrease in survival probability and survival probabilities were non-linearly associated with inbreeding (logistic regression, r^2^ = 0.506, *P* = 5.53 x 10^-7^) (95% CI: 50.4% – 89.5%; Fig. 3D). We estimated the mean number of haploid lethal equivalents to be 4.31 within each adult individual (95% CI: 2.63 – 6.00). Despite the association between *F_ROH_* and fitness, we found no significant differences in realized load between embryonic lethal and mature adult individuals for both loss-of-function (*P* = 0.82, ANOVA) and missense variants (*P* = 0.064, Welch’s ANOVA) (Fig. 3 E, F).

### Genome-wide association mapping of embryonic lethality

The lack of overall increased realized load among embryonic lethal individuals relative to mature adults despite the strong association between *Froh* and survival suggested that rather than a polygenic genetic architecture to inbreeding depression, a large effect recessive lethal could instead be responsible. We used genome-wide association to identify candidate loci associated with embryonic lethality at 5 dpf using SnpSift CaseControl (52) with a recessive genetic model to identify genotypes that were exclusively homozygous derived in embryonic lethal individuals and either homozygous ancestral or heterozygous in mature adults. We found a single region of the genome that surpassed our Bonferroni-corrected significance threshold (*P* < 8.29 x 10^-7^) (Fig. 4).

The candidate region on chromosome 6 encompasses 11 SNPs that span an ∼ 8 Mb region and is associated with 50% of embryonic lethal deaths (N = 14/28). The most significant SNP at Chr6:26048859 is found in an intron of MIB1, an E3 ubiquitin ligase. Mutations in MIB1 cause left ventricular noncompaction cardiomyopathy, characterized by reduced systolic function and pericardial distension (53), consistent with the reduced heart rate of embryonic lethal individuals (Fig. 3C). Remarkably, transgenic zebrafish larva microinjected with MIB1 mutant mRNAs display a straight heart tube, reminiscent of the “stringy” heart phenotype in Devils Hole pupfish (53). All individuals (14/14) that had homozygous derived genotypes at this locus were embryonic lethal.

Along this recessive lethal haplotype, only one SNP was found in a coding region at Chr6:26353888, which introduces a missense mutation in MMP16. MMP16 is a matrix metalloproteinase, a family of proteins crucial for regulating embryonic development in zebrafish (54) and atrial remodeling in atrial fibrillation (55–58). 13/13 individuals that had homozygous derived genotypes at this locus were embryonic lethal. These were the same embryonic lethal individuals with homozygous derived genotypes at MIB1, in addition to a 14^th^ individual with a missing genotype for MMP16.

These two variants along chromosome 6 are unique to Devils Hole pupfish and found in no other desert pupfishes in our dataset. In addition, these variants were not present in our two wild-derived embryonic lethal samples. Overall, this candidate recessive lethal haplotype was segregating at ∼25% in the wild and ∼20% in the refuge. In our two relatively high-quality historical genomes, we found no derived alleles across the 11 candidate lethal variants within this region. Based on the large sizes of the ROHs among the lethal individuals that are homozygous for this recessive lethal haplotype (mean size = 19.3 Mb), we estimate that the inbreeding that generated this homozygous derived haplotype to have occurred within the previous generation.

Finally, on chromosome 8 we found a single hit above our suggestive significance threshold (*P* < 1 x 10^-5^) in an intron of CCBL1, an enzyme implicated in atherosclerotic disease in humans (59). This variant is restricted to the refuge population of Devils Hole pupfish and was not found in wild adults nor wild-derived embryos; 14/15 individuals with a homozygous derived genotype were embryonic lethal. Among the 14 embryonic lethal at this locus, 9 of them overlapped with the individuals with putatively lethal variants on chromosome 6 (Fig. 4).

## Discussion

We investigated the genomic and fitness consequences of long-term small population size in the Devils Hole pupfish by leveraging comparative genomics across spatial, temporal, and ontogenetic scales. Consistent with thousands of generations of isolation at small size, we found Devils Hole pupfish to have one of the lowest genetic diversities ever recorded in the wild. Relative to other populations, Devils Hole pupfish also showed significant shifts in the composition and quantity of genetic load reflecting extensive drift, with most load realized and fixed and with elevated missense load. Despite low inbreeding load, we found evidence of severe inbreeding depression and identified a recessive lethal haplotype with mutations in cardiac development genes MIB1 and MMP16, consistent with the observed “stringy heart” phenotype and reduced heart rate in embryos that died during development. Even at extremely low levels of genetic diversity after thousands of generations of isolation at small population size, our results show that inbreeding still contributes substantially to individual viability and fitness.

Some studies have interpreted evidence of purging and low inbreeding load in small populations as explanations for long-term persistence, suggesting inbreeding depression is unlikely to pose a serious threat (20, 21). By comprehensively linking a recessive lethal haplotype with a specific inbreeding depression phenotype and embryonic lethality, our study demonstrates that long-term small population size can nevertheless impose severe fitness consequences and extinction risk due to high drift load and large effect recessive lethal alleles.

### Demographic history of *C. diabolis*

While we were able to resolve the historical demography of other desert pupfish species, our inference for Devils Hole pupfish lacked resolution beyond ∼3,000 years in the past. Due to long-term low effective population size and recent extreme bottlenecks, most extant genetic variation likely coalesced in a very recent common ancestor, preventing inferences beyond this point. Given a long-term effective population size of 426, the mean coalescent time is only 1,704 (4*N_e_* generations) or approximately 1,278 years in the past based on a generation time of 0.75 years, resulting in lack of power for PSMC beyond this point. Recent studies of other long-term small populations have similarly had trouble inferring deeper demographic history using coalescent-based approaches (46, 60).

Our modern PSMC results are consistent with a more recent colonization of Devils Hole on the order of a few thousand years (34, 61–63) but given aforementioned limitations, it is unclear if divergence occurred earlier. However, our historical PSMC results based on pre-bottleneck genomes that may include signals of deeper coalescent events suggest a decline in effective population size ∼10,000 years ago. This deeper divergence time is consistent with a recalibrated estimate from a previous time-calibrated phylogeny (34) using our recent mutation rate estimate (38), placing the split at 12,750 years (95% CI: 5,250 – 20,400) (Supp. Fig. 4), and supports earlier hypotheses that Devils Hole pupfish became isolated around 10 – 20 kya coinciding with the desiccation of Death Valley towards the late Pleistocene (28).

### Lack of recent decline in genetic diversity for *C. diabolis*

Despite recent bottlenecks, we found no evidence of reduced genetic diversity between the historical (1941) and modern Devils Hole pupfish population (2019-2020), suggesting genetic diversity has consistently been low (Fig. 2B). Several estimates prior to biannual census counts in 1972 suggest the population size was 150 – 300 in the 1950s (64, 65). Furthermore, we know that at least 203 individuals were removed from Devils Hole between 1930 – 1941 to be preserved as specimens, and that hundreds more were translocated out of Devil Hole into nearby springs from 1946-1951, constituting a large proportion of the population removed (66). Notably, substantial inbreeding was found as early as 1941 (Fig. 3A), further supporting our findings of persistently low genetic diversity in the past.

### Genetic architecture of inbreeding depression

Small populations with low inbreeding load are expected to have low risk of inbreeding depression (67–69) and consistent with this expectation, we did not find significantly increased realized load in embryonic lethal individuals relative to adults (Fig. 3E,F). However, this expectation of minimal inbreeding depression is predicated on a relatively polygenic genetic architecture underlying inbreeding depression. Instead, we found that a single recessive lethal haplotype may drive the majority of inbreeding depression in Devils Hole pupfish. This result is unsurprising, given that recessive lethal alleles of large effect have been shown to contribute substantially to inbreeding depression in *Drosophila* (Simmons and Crow 1977; Charlesworth and Charlesworth 1987) and plants (72–74), and are expected to contribute towards inbreeding depression at earlier stages of development than mildly deleterious alleles (75). In addition, high drift load may depress mean population fitness to a sufficient degree that inbreeding depression could be driven by only a few additional deleterious loci that push the burden of genetic load beyond a viable threshold.

### Persistence of a segregating recessive lethal haplotype

Recessive lethals are expected to be rapidly purged in small populations (76–78) or following bottlenecks (79). However, the effectiveness of purging is limited by effective population size and dependent on demographic history (80, 81). In a population as small as Devils Hole pupfish, drift can overpower purging, which may explain why the identified recessive lethal haplotype is still segregating in the population at ∼20%. Additional mechanisms that could underlie the persistence of this recessive lethal haplotype include reproductive compensation (82) and suppression of recombination by inversions (83).

The two candidate lethal variants within this recessive lethal haplotype are unique to Devils Hole pupfish and thus likely arose following initial colonization. The lack of the recessive lethal haplotype in our limited historical genomes suggests that the haplotype was rare before rising in frequency due to bottlenecks in the wild and the founding of the captive refuge population by 29 wild-derived eggs in 2013. Recent studies have identified recessive lethal alleles in bottlenecked populations of Hawaiian crow (84) and Eurasian lynx (85), suggesting the elevation of recessive lethal allele frequencies following bottlenecks may be a generalizable pattern.

### Future of the Devils Hole pupfish

The high levels of drift load observed within Devils Hole pupfish likely depress mean population fitness substantially, as evidenced by the low fecundity, egg viability, and juvenile survivorship observed in the population (27, 86). One potential action to ameliorate drift load is genetic rescue. However, a previous study found that the invasion of only three *C. nevadensis* individuals into a refuge population of Devils Hole pupfish at Point of Rocks rapidly and significantly shifted the gene pool towards *C. nevadensis* alleles in just a few years (87), suggesting that genomic swamping may be a risk. Moving forward, simulating future recessive lethal haplotype frequencies and extinction risk will improve our ability to make informed decisions (21, 88–90) that give Devils Hole pupfish the best chance at continued long-term survival in the wild.

Our study also underscores a new critical value of captive refuges for endangered species: they enable observation of early developmental stages, revealing manifestations of inbreeding depression that would otherwise be virtually undetectable in the wild. Moving forward, genotyping individuals to avoid carrier *x* carrier matings and select non-carriers for reintroduction into the wild will be key for minimizing inbreeding depression and tracking haplotype frequencies over time. To minimize any negative genetic effects from management actions, captive individuals carrying the CCBL1 intronic variant should not be released into the wild because the full effects of this variant on fitness remain unclear.

## Methods

### Sample acquisition

In total, we sequenced 186 new whole genomes including 144 modern samples from fin clips (*C. diabolis* = 41, *C. nevadensis amargosae* = 18, *C. nevadensis mionectes* = 12, *C. nevadensis nevadensis* = 9, *C. nevadensis pectoralis* = 7, *C. nevadensis shoshone* = 11, *C. salinus salinus* = 18, *C. salinus milleri* = 6, *C. radiosus* = 12, *C. macularius* = 10), 28 *C. diabolis* whole lethal embryos (wild = 2, captive = 26) and 14 historical samples from a portion of the caudal peduncle (*C. diabolis* = 9, *C. nevadensis mionectes* = 3, *C. salinus salinus* = 2). Modern samples were collected in the mid-1990s (42, 43), mid-2010s (91–93), and *diabolis* fin clips from 2019-2020 were provided by Ash Meadows Fish Conservation Facility. During downstream SNP filtering, the two historical *C. salinus* individuals were removed due to high missingness (>0.99). In our analyses, we also included 30 previously published Death Valley pupfish genomes on NCBI (PRJNA887195), resulting in a final set of 214 individuals. All sample metadata may be found in Supp. Table 6.

### Extractions

#### RNA

We sampled tissues for brain, eye, and liver for both *C. diabolis* and *C. nevadensis mionectes*, with an additional gill tissue sample for the latter. Each individual was euthanized and immediately placed in dry ice for storage and shipping to UC Berkeley. Tissues were then dissected and placed in RNAlater for one day at 4°C and then stored at −80°C until RNA extractions. We extracted RNA using the Qiagen RNeasy minikit following the manufacturer’s instructions. Library preparation and sequencing was performed by the QB3-Berkeley Genomics core labs (QB3 Genomics, UC Berkeley, Berkeley, CA, RRID:SCR_022170). RNA quality was assessed with an Agilent Fragment Analyzer and libraries were prepped using the KAPA mRNA Hyper Prep kit for extractions with RIN > 6.5. Tissue-specific libraries were sequenced on an Illumina Novaseq 6000 150 PE S4 flowcell, targeting at least 25M reads per sample.

#### Modern WGS

For modern samples, we extracted DNA from fin clips for adults and embryos for embryonic lethal samples with the Qiagen Dneasy Blood & Tissue kit. Library preparation and sequencing was performed by the QB3-Berkeley Genomics core labs (QB3 Genomics, UC Berkeley, Berkeley, CA, RRID:SCR_022170). DNA quality was assessed with an Agilent Fragment Analyzer and libraries were prepped using the KAPA HyperPrep kit for DNA. Libraries were sequenced on an Illumina NovaSeq 6000 150PE S4 flowcell, generating 20Gb minimum of data per sample.

#### Historical WGS

For historical specimen sampling, a small chunk (2mm x 2mm) of the caudal peduncle was dissected. A guanidine DNA extraction (94) and a single-stranded “spotlight” library prep (95) were done at the UCSC Paleogenomics Lab. Proportions of endogenous DNA per sample varied (range = 0.293 – 0.967, mean = 0.80). 2 x 100 PE sequencing was performed at the Duke Sequencing and Genomic Technologies core facility across 4 lanes of a Novaseq X Plus. One historical sample was extracted using a QIAamp DNA FFPE Advanced Kit and sequenced in house at QB3 Berkeley. To repair formalin-induced damage, this one sample was treated with NEBNext FFPE DNA Repair Mix (M6630L, New England Biolabs). A magnetic bead cleanup was performed at 3.0x ratio using Kapa Pure beads and the final product was eluted in 25 µL Buffer EB (10 mM Tris). This in-house sample was sequenced on an Illumina NovaSeq6000 150PE S4 flowcell, generating 10 Gb of data.

#### Reference genome

For each reference genome, high molecular weight extractions were performed at QB3 Berkeley from the tissues stored at −80°C after flash freezing on dry ice. We used PacBio HiFi reads on a single cell of PacBio Sequel II for each sample at QB3 Genomics (UC Berkeley). Omni-C libraries were prepped at the UCSC Paleogenomics Lab and sequenced to 30X coverage by Fulgent Genetics on Illumina Hiseq 2500 150 PE.

### *De novo* genome assembly pipeline

We assembled the genomes with Hifiasm (v0.16.1) (96) using the Savio computational cluster resource at UC Berkeley and masked bacterial and viral contamination based on RefSeq databases. We scaffolded the genomes with Omni-C reads using SALSA (97) and 3DDNA (98). Visualization and manual curation of assemblies was performed with Juicer in Juicebox (v1.11.08) (99). We trained RepeatModeler v2.0.3 (100) on ancestral *Danio rerio* repeats to softmask both generic and clade specific repeats with RepeatMasker v.4.1.2 (101 RepeatMasker Open-4.0). We closed gaps with TGS-GapCloser v1.1.1 (102). We assessed final assembly quality using N50 values, QV scores, and BUSCO scores. We determined genome size and N50 using QUAST v.5.0.2 (103), base-level accuracy (QV score) using Merqury v.1.3 (104), and genome completeness based on the presence of single-copy orthologs present in *Actinopterygii* using BUSCO v.5.2.2 (33).

### Annotation

We annotated the genome with both RNA-seq data and orthology-based RNA-seq reads. For each species-specific tissue sample we cleaned and trimmed reads with fastp v0.23.4 (105) using default parameters and mapped to the corresponding species-specific reference genome using STAR (v2.7.10a) (106) using default parameters, producing a sorted and indexed bam alignment file. Our reference protein fasta was made up of the odb10 vertebrate protein database downloaded from OrthoDB (107) and the protein fasta from the Danio rerio (GRCz11) assembly. For the *C. diabolis* annotation, we used the BRAKER2 (108) pipeline and followed recommendations by running the pipeline separately for RNAseq alignments and reference proteins. The GTF output files from these two runs were combined using TSEBRA, producing a combined gtf output file (109). The agat_sp_extract_sequences.pl function from AGAT was used to extract a protein fasta from the output combined gtf file. In addition, we submitted our protein fasta to the EGGNOG-Mapper web interface for functional annotation (110). Finally, using a custom python script we merged our BRAKER and EGGNOG results. We also used Interproscan v5.52.86 (111) as an independent functional annotation approach.

For the *C. nevadensis mionectes* annotation, we took a slightly different approach. The GeneMark-ES and ProtHint steps of Braker2 failed and only predicted 1490 genes, likely due to tandem repeats with high GC content and a long period that were not captured by RepeatModeler and Repeatmasker; thus GeneMark-ES is likely converging to predicting these repeats rather than real genes (Tomáš Brůna, personal comm.). Similar issues were found in *X. tropicalis* (108). As a workaround, we used GALBA (112) for this step of the pipeline and combined the GALBA output with Braker2 RNAseq output with TSEBRA. The remaining portions of the approach outlined above were identical.

### Genotyping

For modern samples, reads were trimmed with fastp v0.23.4 (105). Reads were mapped to both reference genomes using the following approach. Paired end reads were mapped with BWA mem v0.7.17 (113) using default parameters and duplicates were marked with MarkDuplicatesSpark from GATK v4.5.0.0. For historical samples, reads were trimmed and merged with fastp v0.23.4 (105). Mapping against both reference genomes occurred, as follows. Merged and unmerged reads were mapped using “BWA mem -k=19 r=2.5” following the recommendations of (114), which performs similarly to the standard BWA aln method of (-*I* 1024 -*n* 0.03) used in ancient DNA studies (115). Bam alignment files of unmerged reads were duplicate marked with MarkDuplicatesSpark, while this script (https://github.com/pontussk/samremovedup/samremovedup.py) was used to remove duplicates from merged read bam files. Properly paired reads were extracted from merged read bam files using Samtools v.1.15.1 (116). To assess whether merged read and unmerged properly paired bam alignment files were similar in quality and whether historical samples clustered with modern samples, we plotted each for all historical samples on a PCA with the pcangsd command from angsd v0.94 (117). We also assessed potential DNA damage with Mapdamage v2.2.2 (118). Generally, evidence of C-T substitutions at the 5’ ends of reads and G-A substitutions at 3’ ends for most samples was minimal. Merged and unmerged properly paired bam alignment files were merged with Samtools v.1.15.1 (116). Samples with abnormal base frequency, substitution patterns, or PCA outliers were excluded from downstream analyses. Coverage and mapping quality for all samples was assessed with qualimap v2.2.2 (119) and samtools flagstat v.1.15.1 (120).

### Variant calling

We called variants for each individual with GATK Haplotype Caller (-ERC GVCF) v4.5.0.0 (121). Single-sample GVCFs were imported into a GenomicsDB with GATK GenomicsDBImport for each chromosome and the remaining small scaffolds prior to joint genotyping with GATK GenotypeGVCFs to generate an allsites VCF. Variant sites and invariant sites were separated and missing genotypes were reset to ./. from 0/0 due to how GATK represents missing genotypes in version 4.5.0.0 (122). Variant sites were filtered with GATK recommended hard filters (QD < 2, QUAL < 30, SOR > 3, FS > 60, MQ < 40, MQRankSum < −12.5, ReadPosRankSum < −8) and for biallelic SNPs (123, 124). A genmap mappability mask (-k 150 -E 2) for regions of the genome that uniquely map (mappability > 0.5) was applied. Furthermore, we filtered SNPs to have minimum FMT/GQ of 20, a minimum read depth FMT/DP of 3, and site-level depth INFO/DP of between 1/3X and 2X the mean for minimums and maximums, respectively. We also applied filters for a maximum site missingness of 25% and a minimum minor allele frequency of 0.03. Two historical *C. salinus* individuals were removed due to high missingness (LACM_25258 and LACM_25264). In the end, we ended up with a final high quality VCF of 11,199,716 SNPs and 11,147,432 SNPs for the DHP and MIO references, respectively. For the following analyses, unless otherwise noted, we refer to the VCF aligned against the DHP reference genome.

### Historical demography

To infer long-term historical demography for desert pupfishes, we used the PSMC model (v0.6.5) (37). Briefly, this method infers historical effective population size changes of a given species based on the distribution of time to most recent common ancestor (TMRCA), estimated from the density and distribution of variant sites. We ran PSMC on the highest coverage individual representative of a given species. We excluded from our analyses three outgroups with only a single low coverage sample (*albivelis, fontinalis, eremus*) since at least 18X coverage is recommended (125). For all species, we assumed a mutation rate of 8.09 x 10^-9^ per bp per generation (38) and assumed a generation time of 0.75 years based on Devils Hole pupfish (27). These mutation rates and generation times were used to convert PSMC output into time in years and effective population size, which we plotted over the course of the past 100,000 years.

For PSMC, we called variants on bam files with the *mpileup* (-C 50) and *call* (-c -) commands in bcftools (126) and generated input files using *fq2psmcfa* (-q 20 -s 100). Since we are applying PSMC to individual samples, we used the bcftools pipeline for SNP calling, instead of our GATK called VCF, because bcftools uses single individuals for SNP calling and does not incorporate population allele frequency or Hardy-Weinberg equilibrium into genotype calls. This approach also facilitates comparison to other studies as it has become the standard (125). We ran our inference with the recommended parameters (-N25 -t15 -r5) and default time pattern (4+25*2+4+6) with 100 bootstraps. For Devils Hole pupfish, there was evidence of model overfitting due to insufficient recombination events (<10) for later time segments of the model. As a result, we tried running PSMC with fewer time parameters (4+20*2+6) but there was no improvement.

We applied to PSMC to 2 *diabolis* and 1 *mionectes* relatively high coverage (14 – 15X) historical genomes as well. While inferences of effective population size on recent timescales were stochastic and extremely high likely due to degradation, deeper time inferences of historical and modern *mionectes* matched in overall PSMC trajectory shape.

### Historical admixture

We pruned SNPs in strong linkage disequilibrium using the pruning function (--indep-pairwise 50 5 0.2 in PLINK (v1.9) (127), which left 589,742 SNPs across our 214 individuals. We calculated f-statistics and used the find_graph function with ADMIXTOOLS2 (v2.0.6) (128) in R to infer an admixture graph through fully automated graph exploration. This approach starts with a random admixture graph and improves upon it by making random and targeted changes to the graph, with the outgroup and number of admixture edges set as constants. We started off with automatic graph optimization allowing for no admixture, to establish a baseline. We then tested varying numbers of admixture events (0–10) to see how much admixture resulted in the best likelihood score in terms of represented the data. We ran 5 trials for each number of admixture events given the random start of the process. We used the qpGraph function to calculate out-of-sample scores to account for increasing degrees of freedom given additional admixture events and computed the fit of each graph using bootstrap-resampled SNP blocks to test whether different graphs improved fit to the data. Based on our out-of-sample scores, eight admixture events were inferred to be the best fit for our data. At this level, additional admixture events do not reduce out-of-sample scores. We show the admixture graph with the best scores out of five trials for eight admixture events in Supp. Fig. 1.

### Contemporary population structure

We pruned SNPs in strong linkage disequilibrium using the pruning function (--indep-pairwise 50 10 0.2 in PLINK (v1.9) (127), which left 683,969 SNPs. We then characterized population structure following two complementary approaches. First, we used PLINK to conduct principal component analysis using the –pca function. Afterwards, we used ADMIXTURE (v1.3.0) (129) to assign individuals to varying numbers of population clusters (K = 1-15). We selected K = 10 as the value that best assigned individuals to population clusters based on when cross-validation errors began to level off.

### Genetic diversity

We calculated per-site heterozygosity in non-overlapping sliding windows of 1Mb across the genome in each individual. First, we generated an allsites VCF and extracted invariant sites, which were filtered with a genmap mappability mask (-k 150 -E 2) for regions of the genome that uniquely map (mappability > 0.5), along with RGQ >= 20 and FMT/DP >= 3. This filtered invariant VCF was merged with our filtered variant VCF above as input. We defined heterozygosity as the number of heterozygous genotypes divided by the total number of called genotypes in each window. Windows where fewer than 50% of sites were genotyped were excluded.

### Effective population size

We estimated contemporary effective population size with *currentNe2* (39), which relies on linkage-disequilibrium between markers. Given that estimates are only relevant for a given time point, we restricted our analyses to populations with at least 5 sampled individuals at a given time point. We set the number of chromosomes to 24 and increased the “MAXLOCI” parameter to allow for more markers to be used.

### Inbreeding

We identified runs of homozygosity (ROHs) in our Devils Hole pupfish samples using the BCFtools/ROH command with the Genotypes option (v1.16) (130). Population allele frequency information is necessary as input, but relatedness among Devils Hole pupfish samples could bias ROH calls. As a result, we followed the approach of Leon-Apodaca et al. (131) and used the – make-king option in PLINK (127) to generate a kinship matrix and then the script calc_blue_freq.py (https://github.com/szpiech/blue_af) to generate unbiased allele frequencies estimates by calculating the best linear unbiased estimator for allele frequencies (132), computed by pooling all Devil Hole pupfish samples together. Any negative values were set to zero. These modified allele frequencies were then passed to BCFtools/ROH. We calculated individual inbreeding coefficients in terms of F_ROH_, the proportion of the genome made up of ROHs which we calculated as the sum of ROHs greater than 100 kb divided by the sum of the lengths of the autosomes (1,112,767,855 bp).

We restricted our analyses to samples with at least 15X coverage, given that low coverage can bias ROH estimates with higher rates of both false positives and false negatives (133). We compared levels of inbreeding between wild vs. captive adults and captive adults vs. captive embryonic lethals with two-sample t-tests. We also classified ROHs into various lengths of 0.1 – 1 Mb, 1-5 Mb, 5 – 15 Mb, and greater than 15 Mb and calculated the proportion of the genome that these classes of ROHs comprised in each individual. The age of ROHs were estimated as g = 100 / 2*ROH length (cM), where g is the number of generations to the most recent common ancestor (134). We assumed a generation time of 1 year (27), and an average recombination rate of 3.24 cM/Mb based on sticklebacks (135).

### Inbreeding depression

We modeled the effects of inbreeding depression on survival with a generalized linear model, using RMS (136). In our survival data, mature adults were coded as 1s while embryonic lethal individuals were coded as 0s. For our model, we used a binomial error distribution and logit link with survival as the response variable and F_ROH_ and population modeled as fixed effects. The best supported model was survival ∼ F_ROH_ + population with AIC = 58.7, compared to an alternative model (survival ∼ F_ROH_ * population) with AIC = 60.7. We assessed fit in DHARma (137) and found residuals to be consistent with a uniform distribution, with no evidence of overdispersion or outliers. Quantile deviations were detected, but not significant (Supp. Fig. 4).

To compare the inbreeding depression levels observed in our study to others, we estimated the number of lethal equivalents, defined as a group of deleterious alleles that cause one death on average when made homozygous (138). Following the suggestions of Nietlisbach et al. (139), we refitted our survival model with Poisson errors and a logarithmic link since this approach provides unbiased estimates for binary data. After regressing individual survival on inbreeding coefficients, the negative slope on the logarithmic scale is our number of haploid lethal equivalents. We estimated lethal equivalent confidence intervals with the recommended sandwich estimator (140). We note that the strength of inbreeding depression may vary significantly across life stages (8) and thus we may be under or over estimating the extent of inbreeding depression.

### Genetic load

#### Relative differences in quantity and composition of load

To identify putatively deleterious variants, genotypes in coding sequences were annotated with SnpEff (52). We created a custom SnpEff database using the “build” command and annotated our VCF with the “ann” command. We identified putative loss-of-function SNPs (gained stop codon, lost start codon, lost stop codon) and missense SNPs for our genetic load tally. Next, we polarized alleles based on the genotypes of the *salinus*, *radiosus*, *macularius*, *albivelis*, *eremus*, and *fontinalis* samples (n = 47), which are outgroups relative to *nevadensis* and *diabolis*, given that our SNPs were aligned against the reference genome of a focal species. We restricted our analyses to SNPs where missing genotypes were found in a maximum of 15 outgroup individuals and for which at least 90% of outgroup individuals were in agreement on the same allele, which left 3,103,615 SNPs. The majority allele (> 90%) was inferred to be ancestral, and genotypes were recoded based on their ancestral state (0 = ancestral, 1 = derived). Finally, mutation load estimates were obtained for *diabolis, amargosae*, *mionectes*, *nevadensis*, *pectoralis*, and *shoshone* individuals with less than 30% missing calls using SnpSift (52).

We calculated the masked load (also known as inbreeding load) as the number of putatively deleterious mutations with heterozygous derived alleles in each individual. For the realized load, we calculated both the segregating (mutations that are variable within a population) and drift load (mutations that are fixed within a population). Segregating load was the number of putatively deleterious variants that were homozygous for derived alleles. We calculated drift load as the number of putatively deleterious alleles that were fixed across all the individuals within each respective population. We calculated total load as the total number of derived loss-of-function or nonsynonymous alleles and assessed whether genetic load differed across load types among species using an ANOVA and Tukey’s HSD test to test for pairwise differences. We note that this approach does not quantify the true absolute load harbored by each individual and population given that it is limited to variants with high confidence ancestral allele calls, but nevertheless our approach infers relative differences between species. Finally, we compared realized load among wild, captive, and embryonic lethal *diabolis* populations by summing the number of derived deleterious alleles per individual and normalizing by the number of called deleterious genotypes, to account for differences among samples in coverage and missingness.

#### Fixed SNPs and deletions

We searched for unique fixed variants with predicted functional impacts in Devils Hole pupfish. From our VCF aligned against the *mionectes* reference, we calculated allele frequencies among all of our Devils Hole pupfish samples and among all other pupfishes to identify variants that were variant in Devils Hole pupfish samples but invariant in all other samples. True fixed variants would not have been found if we had used *diabolis* reference, since that would have required the reference individual to be invariant. We further narrowed down our list of variants by intersecting the variants with exons, looking only at fixed SNPs (AF=1), and identifying which SNPs were predicted to be loss-of-function based on Snpeff (52).

We also searched for unique deletions within Devils Hole pupfish by calling, merging, genotyping, and filtering structural variants with DELLY (v1.1.6, (141)). We used as input bam files aligned against the *C. nevadensis mionectes* reference. We intersected deletion calls with our annotation in order to restrict our analyses to deletions that intersected with exons. Once identified, we searched for the putative deletions in the genomes of all other desert pupfish species. We visualized deletions to confirm their presence with WALLY (v0.5.8) (142), given that calling of structural variants often requires manual curation in the absence of long-reads (143).

### Heart rate

We measured the heart rate of healthy and embryonic lethal embryos using 1 minute timed videos. For each video of development, we counted beats three times with a clicker and took the average number of beats across three trials divided by the length of the video as the heart rate. We compared heart rates between healthy and embryonic lethal embryos with a two-sample *t*-test.

### Genome-wide association mapping of embryonic lethality

We tested for SNPs associated with recessive embryonic lethality using the “CaseControl” function in SnpSift, which can perform association testing under a recessive genetic model by contrasting homozygous genotypes for one set of alleles against other genotypes (heterozygotes and homozygotes of the opposite set). For all SNPs within *C. diabolis*, we performed association testing using Fisher’s exact tests (2×2 contingency tables), comparing the frequency of homozygous genotypes (0/0 or 1/1) between wild and captive embryos that died at 5-6 dpf (n=28) and mature adults (n=41). We checked genotypes in outgroups to infer whether alleles were ancestral or derived. We plotted −log10(*p*-values) with a suggestive p-value threshold of *P* < 1 x 10^-5^ and a Bonferroni significance threshold of *P* < 8.29 x 10^-7^ based on 0.05 / 60,333 (number of SNPs). We did not control for relatedness nor population structure among our samples because our goal was to identify candidate recessive lethal variants that are realized upon becoming homozygous, which primarily occurs through shared recent common ancestors. To assess the potential functional consequences of candidate variants underlying the two strongest association signals, we cross-referenced our candidate lethal variants on chromosomes 6 and 8 with our snpEff annotation database to classify SNPs based on their genomic context and predicted effect on protein sequence.

## Data availability

All novel sequence data generated by this study is deposited to NCBI’s SRA database under XXX. *De novo* genome assemblies are publicly available on NCBI at GCA_030533445.1 and GCA_030533455.1.

## Acknowledgements

This research was funded by U.S. Fish and Wildlife, NSF CAREER 1749764, and NIH 5R01DE027052-02 awards to CHM. We thank U.S. Fish and Wildlife for permission to collect and possess samples. The findings and conclusions in this article are those of the authors and do not necessarily represent the views of the U.S. Fish and Wildlife Service. All research procedures and animal care protocols (AUP-2021-02-14062-1 and AUP-2021-07-14515) were approved by the University of California, Berkeley Animal Care and Use committee. D.T. was supported by an NSF GRFP (DGE 1752814), the Museum of Vertebrate Zoology, and a Philomathia Graduate Fellowship in the Environmental Sciences. We thank U.S. Fish & Wildlife and the National Park Service for their work in managing and conserving this species. We thank Craig Miller, Stephanie Carlson, Stephen Gaughran, and members of the Martin lab for valuable feedback and discussion of this project. We thank Dave Catania at California Academy of Sciences, Hernán López-Fernández and Randy Singer at the University of Michigan Museum of Zoology, Todd Clardy and William Ludt at the Natural History Museum of Los Angeles County, Justin Mann at the Tulane University Biodiversity Research Institute, and Chris Feldman at the University of Nevada, Reno, Museum of Natural History for loaning specimens for sequencing (CAS 22994, 82806; UMMZ 134803, 140460; LACM 25258, 25264; TU 75085; UNR 4284, 4285, 4287, 14503). Voucher specimens for all sequenced individuals are deposited in the Museum of Vertebrate Zoology (MVZ:Fish:350,351,7071,7072,3880) and available upon request.

## Author Contributions

D.T., and C.H.M. conceptualized the project. D.T. performed all analyses. D.T., N.S., and V.M. performed lab work. C.H.M. provided funding for sequencing and analysis and acquired all samples. D.T. wrote the manuscript. B.T., S.L., K.W., O.F., M.S., and J.G. collected samples. All authors edited and reviewed the manuscript.

## References

1. M. E. Soulé, What Is Conservation Biology? BioScience 35, 727–734 (1985).

2. B. A. Melbourne, A. Hastings, Extinction risk depends strongly on factors contributing to stochasticity. Nature 454, 100–103 (2008).

3. D. H. Reed, R. Frankham, Correlation between Fitness and Genetic Diversity. Conservation Biology 17, 230–237 (2003).

4. Y. Willi, J. V. Buskirk, A. A. Hoffmann, Limits to the Adaptive Potential of Small Populations. *Annual Review of Ecology*, Evolution, and Systematics 37, 433–458 (2006).

5. M. Kardos, et al., The crucial role of genome-wide genetic variation in conservation. Proceedings of the National Academy of Sciences 118, e2104642118 (2021).

6. R. Lande, Genetics and demography in biological conservation. Science (New York, N.Y.) 241 (1988).

7. L. F. Keller, D. M. Waller, Inbreeding effects in wild populations. Trends in Ecology & Evolution 17, 230–241 (2002).

8. M. A. Stoffel, S. E. Johnston, J. G. Pilkington, J. M. Pemberton, Genetic architecture and lifetime dynamics of inbreeding depression in a wild mammal. Nat Commun 12, 2972 (2021).

9. M. Kardos, et al., Inbreeding depression explains killer whale population dynamics. Nat Ecol Evol 7, 675–686 (2023).

10. A. L. K. Mattila, et al., High genetic load in an old isolated butterfly population. Proceedings of the National Academy of Sciences 109, E2496–E2505 (2012).

11. T. van der Valk, D. Díez-del-Molino, T. Marques-Bonet, K. Guschanski, L. Dalén, Historical Genomes Reveal the Genomic Consequences of Recent Population Decline in Eastern Gorillas. Current Biology 29, 165–170.e6 (2019).

12. C. van Oosterhout, Mutation load is the spectre of species conservation. Nat Ecol Evol 4, 1004–1006 (2020).

13. G. Bertorelle, et al., Genetic load: genomic estimates and applications in non-model animals. Nat Rev Genet 23, 492–503 (2022).

14. M. Lynch, W. Gabriel, Mutation load and the survival of small populations. Evol 44, 1725–1737 (1990).

15. M. Lynch, J. Conery, R. Bürger, Mutational Meltdowns in Sexual Populations. Evolution 49, 1067–1080 (1995).

16. M. Lynch, J. Conery, R. Bürger, Mutation Accumulation and the Extinction of Small Populations. The American Naturalist 146, 489–518 (1995).

17. C. Grossen, F. Guillaume, L. F. Keller, D. Croll, Purging of highly deleterious mutations through severe bottlenecks in Alpine ibex. Nature Communications 11, 1001 (2020).

18. D. Kleinman-Ruiz, et al., Purging of deleterious burden in the endangered Iberian lynx. Proceedings of the National Academy of Sciences 119, e2110614119 (2022).

19. R. J. Laws, I. G. Jamieson, Is lack of evidence of inbreeding depression in a threatened New Zealand robin indicative of reduced genetic load? Animal Conservation 14, 47–55 (2011).

20. J. A. Robinson, C. Brown, B. Y. Kim, K. E. Lohmueller, R. K. Wayne, Purging of Strongly Deleterious Mutations Explains Long-Term Persistence and Absence of Inbreeding Depression in Island Foxes. Current Biology 28, 3487–3494.e4 (2018).

21. J. A. Robinson, et al., The critically endangered vaquita is not doomed to extinction by inbreeding depression. Science 376, 635–639 (2022).

22. A. Khan, et al., Genomic evidence for inbreeding depression and purging of deleterious genetic variation in Indian tigers. PNAS 118 (2021).

23. R. Lande, Risk of population extinction from fixation of new deleterious mutations. Evol 48, 1460–1469 (1994).

24. P. W. Hedrick, A. Garcia-Dorado, Understanding Inbreeding Depression, Purging, and Genetic Rescue. Trends in Ecology & Evolution 31, 940–952 (2016).

25. N. Dussex, H. E. Morales, C. Grossen, L. Dalén, C. van Oosterhout, Purging and accumulation of genetic load in conservation. Trends in Ecology & Evolution 38, 961–969 (2023).

26. S. Mathur, J. M. Tomeček, L. A. Tarango-Arámbula, R. M. Perez, J. A. DeWoody, An evolutionary perspective on genetic load in small, isolated populations as informed by whole genome resequencing and forward-time simulations. Evolution 77, 690–704 (2023).

27. J. E. Deacon, F. R. Taylor, J. W. Pedretti, Egg viability and ecology of Devils Hole pupfish: Insights from captive propagation. The Southwestern Naturalist 40, 216–223 (1995).

28. R. R. Miller, Coevolution of deserts and pupfishes (genus Cyprinodon) in the American Southwest, in Naiman, R. J., and Soltz, D. L., eds, Fishes in North American Deserts (1981).

29. T. K. Lowenstein, et al., 200 k.y. paleoclimate record from Death Valley salt core. Geology 27, 3–6 (1999).

30. D. Tian, A. H. Patton, B. J. Turner, C. H. Martin, Severe inbreeding, increased mutation load and gene loss-of-function in the critically endangered Devils Hole pupfish. Proceedings of the Royal Society B: Biological Sciences 289, 20221561 (2022).

31. Cappaert v. United States (1976).

32. M. E. Andersen, J. E. Deacon, Population Size of Devils Hole Pupfish (Cyprinodon diabolis) Correlates with Water Level. cope 2001, 224–228 (2001).

33. M. Seppey, M. Manni, E. M. Zdobnov, BUSCO: Assessing Genome Assembly and Annotation Completeness. Methods Mol Biol 1962, 227–245 (2019).

34. C. H. Martin, Crawford Jacob E., Turner Bruce J., Simons Lee H., Diabolical survival in Death Valley: recent pupfish colonization, gene flow and genetic assimilation in the smallest species range on earth. Proceedings of the Royal Society B: Biological Sciences 283, 20152334 (2016).

35. I. La Rivers, The Dryopoidea Known or Expected to Occur in the Nevada Area (Coleoptera). Wasmann Journal of Biology 8, 106 (1950).

36. I. La Rivers, T. J. Trelease, An Annotated Check List of the Fishes of Nevada. California Fish and Game 38, 119 (1952).

37. H. Li, R. Durbin, Inference of human population history from individual whole-genome sequences. Nature 475, 493–496 (2011).

38. D. Tian, et al., Estimate of the mutation rate in the endangered Devils Hole pupfish provides support for the drift-barrier hypothesis at an outlying extreme. [Preprint] (2026). Available at: https://www.biorxiv.org/content/10.64898/2026.01.11.698928v1.

39. E. Santiago, C. Köpke, A. Caballero, Accounting for population structure and data quality in demographic inference with linkage disequilibrium methods. Nat Commun 16, 6054 (2025).

40. A. A. Echelle, T. E. Dowling, Mitonchondrial DNA variation and evolution of the Death Valley pupfishes (Cyprinodon, Cyprinodontidae). Evolution 46, 193–206 (1992).

41. A. A. Echelle, A. F. Echelle, Allozyme Perspective on Mitochondrial DNA Variation and Evolution of the Death Valley Pupfishes (Cyprinodontidae: Cyprinodon). Copeia 1993, 275–287 (1993).

42. D. D. Duvernell, B. J. Turner, Evolutionary genetics of Death Valley pupfish populations: mitochondrial DNA sequence variation and population structure. Molecular Ecology 7, 279–288 (1998).

43. D. D. Duvernell, B. J. Turner, Variation and Divergence of Death Valley Pupfish Populations at Retrotransposon-Defined Loci. Molecular Biology and Evolution 16, 363–371 (1999).

44. E. Pister, Species in a Bucket. Nat. Hist. 102, 14- (1993).

45. P. A. Morin, et al., Reference genome and demographic history of the most endangered marine mammal, the vaquita. Molecular Ecology Resources 21, 1008–1020 (2021).

46. M. Gabrielli, et al., The relationship between genomic variation and genetic load: insights from small island populations. [Preprint] (2026). Available at: https://www.biorxiv.org/content/10.64898/2026.03.06.710193v1.

47. M. García, et al., Phosphofructo-1-Kinase Deficiency Leads to a Severe Cardiac and Hematological Disorder in Addition to Skeletal Muscle Glycogenosis. PLOS Genetics 5, e1000615 (2009).

48. S. Tarui, et al., Phosphofructokinase deficiency in skeletal muscle. A new type of glycogenosis. Biochem Biophys Res Commun 19, 517–523 (1965).

49. G. L. McKnight, T. S. Cardillo, F. Sherman, An extensive deletion causing overproduction of yeast iso-2-cytochrome c. Cell 25, 409–419 (1981).

50. P. J. van der Spek, et al., Chromosomal Localization of Three Repair Genes: The Xeroderma Pigmentosum Group C Gene and Two Human Homologs of Yeast RAD23. Genomics 23, 651–658 (1994).

51. M. Grønbæk-Thygesen, C. Kampmeyer, K. Hofmann, R. Hartmann-Petersen, The moonlighting of RAD23 in DNA repair and protein degradation. Biochimica et Biophysica Acta (BBA) - Gene Regulatory Mechanisms 1866, 194925 (2023).

52. P. Cingolani, et al., A program for annotating and predicting the effects of single nucleotide polymorphisms, SnpEff. Fly (Austin) 6, 80–92 (2012).

53. G. Luxán, et al., Mutations in the NOTCH pathway regulator MIB1 cause left ventricular noncompaction cardiomyopathy. Nat Med 19, 193–201 (2013).

54. J. Zhang, S. Bai, X. Zhang, H. Nagase, M. P. Sarras, The expression of novel membrane-type matrix metalloproteinase isoforms is required for normal development of zebrafish embryos. Matrix Biology 22, 279–293 (2003).

55. Y. Nakano, et al., Matrix metalloproteinase-9 contributes to human atrial remodeling during atrial fibrillation. JACC 43, 818–825 (2004).

56. F. G. Spinale, Myocardial Matrix Remodeling and the Matrix Metalloproteinases: Influence on Cardiac Form and Function. Physiological Reviews 87, 1285–1342 (2007).

57. G. W. Moe, et al., Matrix Metalloproteinase Inhibition Attenuates Atrial Remodeling and Vulnerability to Atrial Fibrillation in a Canine Model of Heart Failure. Journal of Cardiac Failure 14, 768–776 (2008).

58. M. Fragão-Marques, et al., Atrial matrix remodeling in atrial fibrillation patients with aortic stenosis. BMC Cardiovasc Disord 20, 468 (2020).

59. R. Baumgartner, et al., Evidence that a deviation in the kynurenine pathway aggravates atherosclerotic disease in humans. Journal of Internal Medicine 289, 53–68 (2021).

60. A. D. Foote, et al., “Type D” killer whale genomes reveal long-term small population size and low genetic diversity. J Hered 114, 94–109 (2023).

61. J. M. Reed, C. A. Stockwell, Evaluating an icon of population persistence: the Devil’s Hole pupfish. Proceedings of the Royal Society B: Biological Sciences 281, 20141648 (2014).

62. C. H. Martin, et al., The complex effects of demographic history on the estimation of substitution rate: concatenated gene analysis results in no more than twofold overestimation. Proceedings of the Royal Society B: Biological Sciences 284, 20170537 (2017).

63. C. H. Martin, S. Höhna, New evidence for the recent divergence of Devil’s Hole pupfish and the plausibility of elevated mutation rates in endangered taxa. Mol Ecol 27, 831–838 (2018).

64. R. R. Miller, Field notes, Box 29A. (1950).

65. R. J. Hoffman, “Chronology of diving activities and underground surveys in Devils Hole and Devils Hole Cave, Nye County, Nevada, 1950-86” (U.S. Geological Survey, 1988).

66. K. Brown C., Devils Hole Pupfish: The Unexpected Survival of an Endangered Species in the Modern American West (2021).

67. N. Pérez-Pereira, et al., Long-term exhaustion of the inbreeding load in Drosophila melanogaster. Heredity 127, 373–383 (2021).

68. C. C. Kyriazis, J. A. Robinson, K. E. Lohmueller, Long runs of homozygosity are reliable genomic markers of inbreeding depression. Trends in Ecology & Evolution 40, 874–884 (2025).

69. T. Bataillon, M. Kirkpatrick, Inbreeding depression due to mildly deleterious mutations in finite populations: size does matter. Genet Res 75, 75–81 (2000).

70. M. J. Simmons, J. F. Crow, Mutations affecting fitness in Drosophila populations. Annu Rev Genet 11, 49–78 (1977).

71. D. Charlesworth, B. Charlesworth, Inbreeding depression and its evolutionary consequences. Annual Review of Ecology, Evolution, and Systematics 18, 237–268 (1987).

72. E. J. Klekowski, Mutational Load in Clonal Plants: A Study of Two Fern Species. Evolution 38, 417–426 (1984).

73. P. W. Hedrick, GENETIC LOAD AND THE MATING SYSTEM IN HOMOSPOROUS FERNS. Evol 41, 1282–1289 (1987).

74. J. H. Willis, Genetic analysis of inbreeding depression caused by chlorophyll-deficient lethals in Mimulus guttatus. Heredity 69, 562–572 (1992).

75. B. C. Husband, D. W. Schemske, Evolution of the magnitude and timing of inbreeding depression in plants. Evolution 50, 54–70 (1996).

76. M. Nei, The frequency distribution of lethal chromosomes in finite populations. Proceedings of the National Academy of Sciences 60, 517–524 (1968).

77. P. W. Hedrick, Purging inbreeding depression and the probability of extinction: full-sib mating. Heredity 73, 363–372 (1994).

78. P. W. Hedrick, Lethals in finite populations. Evolution 56, 654–657 (2002).

79. M. Kirkpatrick, P. Jarne, The Effects of a Bottleneck on Inbreeding Depression and the Genetic Load. The American Naturalist 155, 154–167 (2000).

80. S. Glémin, How are deleterious mutations purged? Drift versus nonrandom mating. Evolution 57, 2678–2687 (2003).

81. J. Robinson, C. C. Kyriazis, S. C. Yuan, K. E. Lohmueller, Deleterious Variation in Natural Populations and Implications for Conservation Genetics. Annu Rev Anim Biosci 11, 93–114 (2023).

82. C. Ober, T. Hyslop, W. W. Hauck, Inbreeding Effects on Fertility in Humans: Evidence for Reproductive Compensation. The American Journal of Human Genetics 64, 225–231 (1999).

83. S. B. Marion, M. A. F. Noor, Interrogating the Roles of Mutation-Selection Balance, Heterozygote Advantage, and Linked Selection in Maintaining Recessive Lethal Variation in Natural Populations. Annu Rev Anim Biosci 11, 77–91 (2023).

84. C. C. Kyriazis, et al., Elevated recessive lethal frequencies drive hatching failure following near extinction in ‘Alalā, the Hawaiian crow. [Preprint] (2026). Available at: https://www.biorxiv.org/content/10.64898/2026.03.24.713968v1.

85. J. Niehaus, et al., Evolutionary dynamics of a lethal recessive allele in reintroduced fragmented lynx populations. [Preprint] (2025). Available at: https://www.biorxiv.org/content/10.1101/2025.11.06.686959v1.

86. J. Gumm, et al., Insights from 10 Years of operations at the Ash Meadows Fish Conservation Facility. Desert Fishes Council 55th Annual Symposium (2023).

87. A. P. Martin, A. A. Echelle, G. Zegers, S. Baker, C. L. Keeler-Foster, Dramatic shifts in the gene pool of a managed population of an endangered species may be exacerbated by high genetic load. Conserv Genet 13, 349–358 (2012).

88. C. C. Kyriazis, J. A. Robinson, K. E. Lohmueller, Using Computational Simulations to Model Deleterious Variation and Genetic Load in Natural Populations. Am Nat 202, 737–752 (2023).

89. J. E. Beaman, et al., A Guide for Developing Demo-Genetic Models to Simulate Genetic Rescue. Evolutionary Applications 18, e70092 (2025).

90. C. C. Kyriazis, et al., Population genomics of recovery and extinction in Hawaiian honeycreepers. Current Biology 35, 2697–2708.e4 (2025).

91. A. A. Del Core, C. S. Cleveland, S. C. Lema, Complete mitochondrial genome of the Salt Creek pupfish, Cyprinodon salinus salinus: characterization and identification of single nucleotide polymorphisms (SNPs). Mitochondrial DNA Part B 6, 2229–2232 (2021).

92. S. C. Lema, M. I. Chow, A. H. Dittman, D. May, M. J. Housh, Accustomed to the heat: Temperature and thyroid hormone influences on oogenesis and gonadal steroidogenesis pathways vary among populations of Amargosa pupfish (*Cyprinodon nevadensis amargosae*). Comparative Biochemistry and Physiology Part A: Molecular & Integrative Physiology 272, 111280 (2022).

93. M. J. Housh, J. Telish, K. L. Forsgren, S. C. Lema, Fluctuating and Stable High Temperatures Differentially Affect Reproductive Endocrinology of Female Pupfish. Integrative Organismal Biology 6 (2024).

94. N. Straube, et al., Successful application of ancient DNA extraction and library construction protocols to museum wet collection specimens. Molecular Ecology Resources 21, 2299–2315 (2021).

95. J. D. Kapp, R. E. Green, B. Shapiro, A Fast and Efficient Single-stranded Genomic Library Preparation Method Optimized for Ancient DNA. Journal of Heredity 112, 241–249 (2021).

96. H. Cheng, G. T. Concepcion, X. Feng, H. Zhang, H. Li, Haplotype-resolved de novo assembly using phased assembly graphs with hifiasm. Nat Methods 18, 170–175 (2021).

97. J. Ghurye, M. Pop, S. Koren, D. Bickhart, C.-S. Chin, Scaffolding of long read assemblies using long range contact information. BMC Genomics 18, 527 (2017).

98. O. Dudchenko, et al., De novo assembly of the Aedes aegypti genome using Hi-C yields chromosome-length scaffolds. Science 356, 92–95 (2017).

99. N. C. Durand, et al., Juicer Provides a One-Click System for Analyzing Loop-Resolution Hi-C Experiments. Cell Syst 3, 95–98 (2016).

100. J. M. Flynn, et al., RepeatModeler2 for automated genomic discovery of transposable element families. Proc Natl Acad Sci U S A 117, 9451–9457 (2020).

101. A. Smit, R. Hubley, P. Green, RepeatMasker Open-4.0.

102. M. Xu, et al., TGS-GapCloser: A fast and accurate gap closer for large genomes with low coverage of error-prone long reads. GigaScience 9, giaa094 (2020).

103. A. Gurevich, V. Saveliev, N. Vyahhi, G. Tesler, QUAST: quality assessment tool for genome assemblies. Bioinformatics 29, 1072–1075 (2013).

104. A. Rhie, B. P. Walenz, S. Koren, A. M. Phillippy, Merqury: reference-free quality, completeness, and phasing assessment for genome assemblies. Genome Biol 21, 245 (2020).

105. S. Chen, Y. Zhou, Y. Chen, J. Gu, fastp: an ultra-fast all-in-one FASTQ preprocessor. Bioinformatics 34, i884–i890 (2018).

106. A. Dobin, et al., STAR: ultrafast universal RNA-seq aligner. Bioinformatics 29, 15–21 (2013).

107. E. V. Kriventseva, et al., OrthoDB v10: sampling the diversity of animal, plant, fungal, protist, bacterial and viral genomes for evolutionary and functional annotations of orthologs. Nucleic Acids Res 47, D807–D811 (2019).

108. T. Brůna, K. J. Hoff, A. Lomsadze, M. Stanke, M. Borodovsky, BRAKER2: automatic eukaryotic genome annotation with GeneMark-EP+ and AUGUSTUS supported by a protein database. NAR Genom Bioinform 3, lqaa108 (2021).

109. L. Gabriel, K. J. Hoff, T. Brůna, M. Borodovsky, M. Stanke, TSEBRA: transcript selector for BRAKER. BMC Bioinformatics 22, 566 (2021).

110. C. P. Cantalapiedra, A. Hernández-Plaza, I. Letunic, P. Bork, J. Huerta-Cepas, eggNOG-mapper v2: Functional Annotation, Orthology Assignments, and Domain Prediction at the Metagenomic Scale. Mol Biol Evol 38, 5825–5829 (2021).

111. P. Jones, et al., InterProScan 5: genome-scale protein function classification. Bioinformatics 30, 1236–1240 (2014).

112. T. Brůna, et al., Galba: genome annotation with miniprot and AUGUSTUS. BMC Bioinformatics 24, 327 (2023).

113. H. Li, Aligning sequence reads, clone sequences and assembly contigs with BWA-MEM. [Preprint] (2013). Available at: http://arxiv.org/abs/1303.3997.

114. W. Xu, et al., An efficient pipeline for ancient DNA mapping and recovery of endogenous ancient DNA from whole-genome sequencing data. Ecology and Evolution 11, 390–401 (2021).

115. M. Schubert, et al., Improving ancient DNA read mapping against modern reference genomes. BMC Genomics 13, 178 (2012).

116. P. Danecek, et al., Twelve years of SAMtools and BCFtools. GigaScience 10, giab008 (2021).

117. J. Meisner, A. Albrechtsen, Inferring Population Structure and Admixture Proportions in Low-Depth NGS Data. Genetics 210, 719–731 (2018).

118. H. Jónsson, A. Ginolhac, M. Schubert, P. L. F. Johnson, L. Orlando, mapDamage2.0: fast approximate Bayesian estimates of ancient DNA damage parameters. Bioinformatics 29, 1682–1684 (2013).

119. K. Okonechnikov, A. Conesa, F. García-Alcalde, Qualimap 2: advanced multi-sample quality control for high-throughput sequencing data. Bioinformatics 32, 292–294 (2016).

120. H. Li, et al., The Sequence Alignment/Map format and SAMtools. Bioinformatics 25, 2078–2079 (2009).

121. A. McKenna, et al., The Genome Analysis Toolkit: A MapReduce framework for analyzing next-generation DNA sequencing data. Genome Research 20, 1297 (2010).

122. D. Caetano-Anolles, GenotypeGVCFs and the death of the dot (obsolete as of GATK 4.6.0.0). GATK (2024). Available at: https://gatk.broadinstitute.org/hc/en-us/articles/6012243429531-GenotypeGVCFs-and-the-death-of-the-dot-obsolete-as-of-GATK-4-6-0-0.

123. M. A. DePristo, et al., A framework for variation discovery and genotyping using next-generation DNA sequencing data. Nat Genet 43, 491–498 (2011).

124. G. A. Van der Auwera, et al., From FastQ Data to High-Confidence Variant Calls: The Genome Analysis Toolkit Best Practices Pipeline. Current Protocols in Bioinformatics 43, 11.10.1-11.10.33 (2013).

125. K. Nadachowska-Brzyska, R. Burri, L. Smeds, H. Ellegren, PSMC analysis of effective population sizes in molecular ecology and its application to black-and-white Ficedula flycatchers. Mol Ecol 25, 1058–1072 (2016).

126. H. Li, A statistical framework for SNP calling, mutation discovery, association mapping and population genetical parameter estimation from sequencing data. Bioinformatics 27, 2987–2993 (2011).

127. C. C. Chang, et al., Second-generation PLINK: rising to the challenge of larger and richer datasets. Gigascience 4, 7 (2015).

128. R. Maier, et al., On the limits of fitting complex models of population history to f-statistics. eLife 12, e85492 (2023).

129. D. H. Alexander, J. Novembre, K. Lange, Fast model-based estimation of ancestry in unrelated individuals. Genome Res 19, 1655–1664 (2009).

130. V. Narasimhan, et al., BCFtools/RoH: a hidden Markov model approach for detecting autozygosity from next-generation sequencing data. Bioinformatics 32, 1749–1751 (2016).

131. A. V. Leon-Apodaca, et al., Genomic Consequences of Isolation and Inbreeding in an Island Dingo Population. Genome Biology and Evolution 16, evae130 (2024).

132. M. S. McPeek, X. Wu, C. Ober, Best linear unbiased allele-frequency estimation in complex pedigrees. Biometrics 60, 359–367 (2004).

133. G. A. A. Silva, A. M. Harder, K. B. Kirksey, S. Mathur, J. R. Willoughby, Detectability of runs of homozygosity is influenced by analysis parameters and population-specific demographic history. PLOS Computational Biology 20, e1012566 (2024).

134. E. A. Thompson, Identity by Descent: Variation in Meiosis, Across Genomes, and in Populations. Genetics 194, 301–326 (2013).

135. V. Venu, et al., Fine-scale contemporary recombination variation and its fitness consequences in adaptively diverging stickleback fish. Nat Ecol Evol 8, 1337–1352 (2024).

136. F. E. Harrell, rms: Regression Modeling Strategies. (2025). Deposited 4 April 2025.

137. F. Hartig, L. Lohse, M. de S. leite, DHARMa: Residual Diagnostics for Hierarchical (Multi-Level / Mixed) Regression Models. (2024). Deposited 18 October 2024.

138. N. E. Morton, J. F. Crow, H. J. Muller, An estimate of the mutational damage in man from data on consanguineous marriages. Proc Natl Acad Sci U S A 42, 855–863 (1956).

139. P. Nietlisbach, S. Muff, J. M. Reid, M. C. Whitlock, L. F. Keller, Nonequivalent lethal equivalents: Models and inbreeding metrics for unbiased estimation of inbreeding load. Evolutionary Applications 12, 266–279 (2019).

140. G. Zou, A Modified Poisson Regression Approach to Prospective Studies with Binary Data. American Journal of Epidemiology 159, 702–706 (2004).

141. T. Rausch, et al., DELLY: structural variant discovery by integrated paired-end and split-read analysis. Bioinformatics 28, i333–i339 (2012).

142. T. Rausch, et al., Long-read sequencing of diagnosis and post-therapy medulloblastoma reveals complex rearrangement patterns and epigenetic signatures. Cell Genomics 3, 100281 (2023).

143. J. A. Hemker, H. R. Gellert, J. A. Smiley-Rhodes, B. Y. Kim, D. A. Petrov, Manual validation finds only ultra-long long-read sequencing enables faithful, population-level structural variant calling in Drosophila melanogaster euchromatin. [Preprint] (2025). Available at: https://www.biorxiv.org/content/10.1101/2025.04.21.649852v1.

