## Supplemental Materials for "Long-term small effective population size, inbreeding, and a recessive lethal haplotype drive premature death in the endangered Devils Hole pupfish (*Cyprinodon diabolis*)"

**This file includes:**

Supplementary Text
Figures S1 – S7
Tables S1 – S6
SI References

### Supplemental Figures

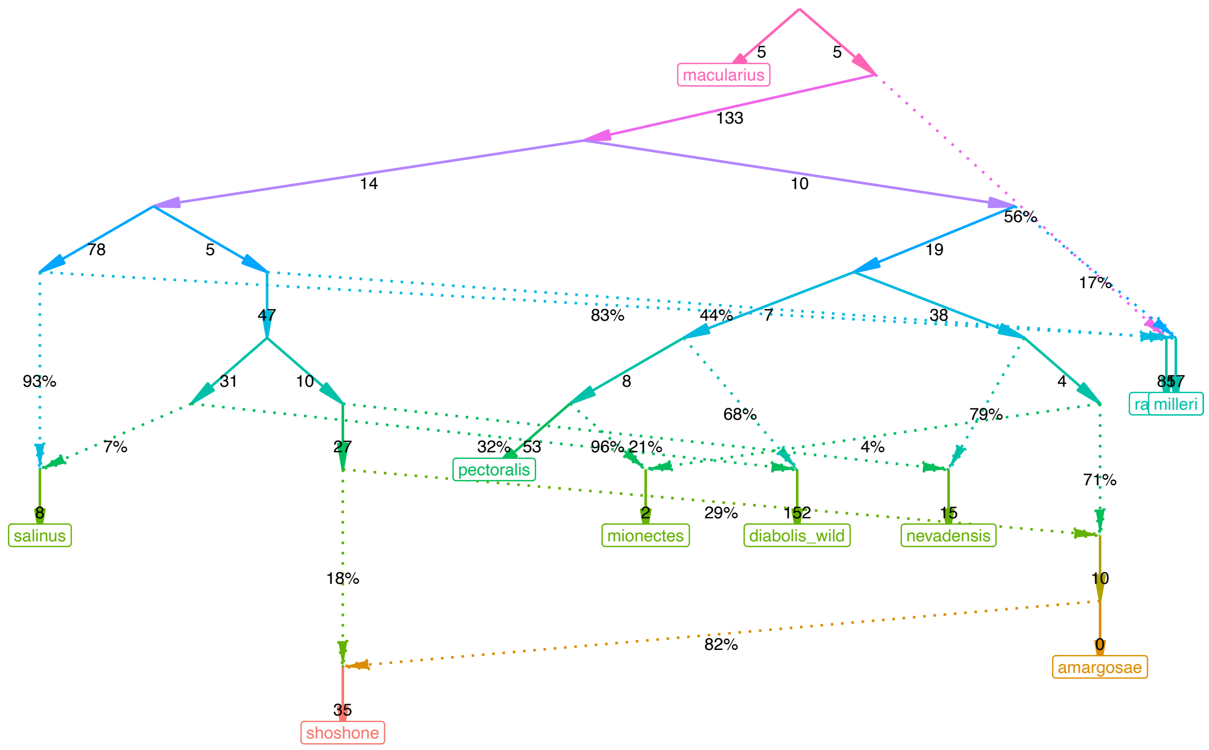


**Supp. Fig. 1**. Inferred admixture graph, using *C. macularius* as an outgroup. All *f*-statisitcs were calculated and plotted using *admixtools2* (Maier et al. 2023). We determined eight to be the optimal number of admixture events and plot the best scoring graph (out-of-sample score 163.6) here.

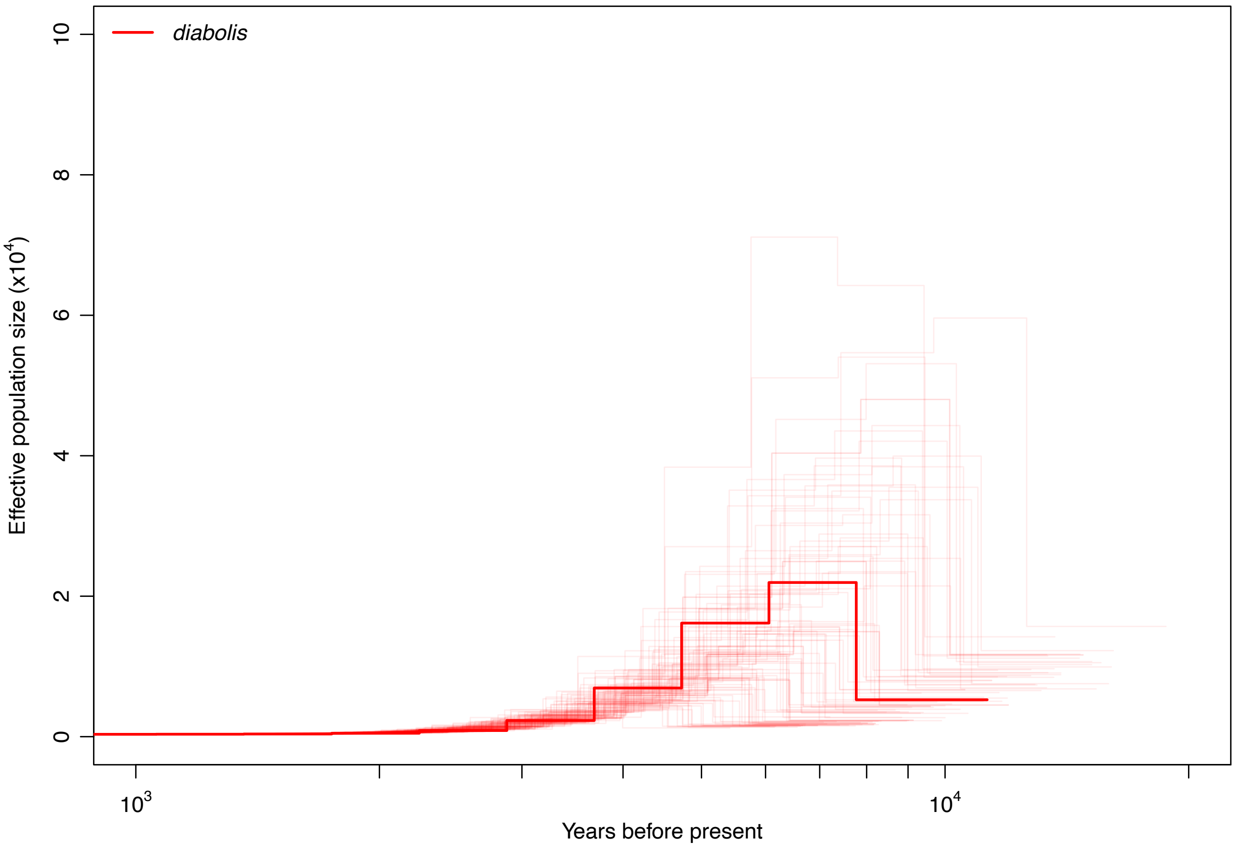


**Supp. Fig. 2**. Zoomed in view of historic effective population size inferred by PSMC for a Devils Hole pupfish individual (RT1, 25X coverage). Wide range of bootstrap estimates illustrates lack of power and increasing uncertainty as the effective population inference goes further back in time.

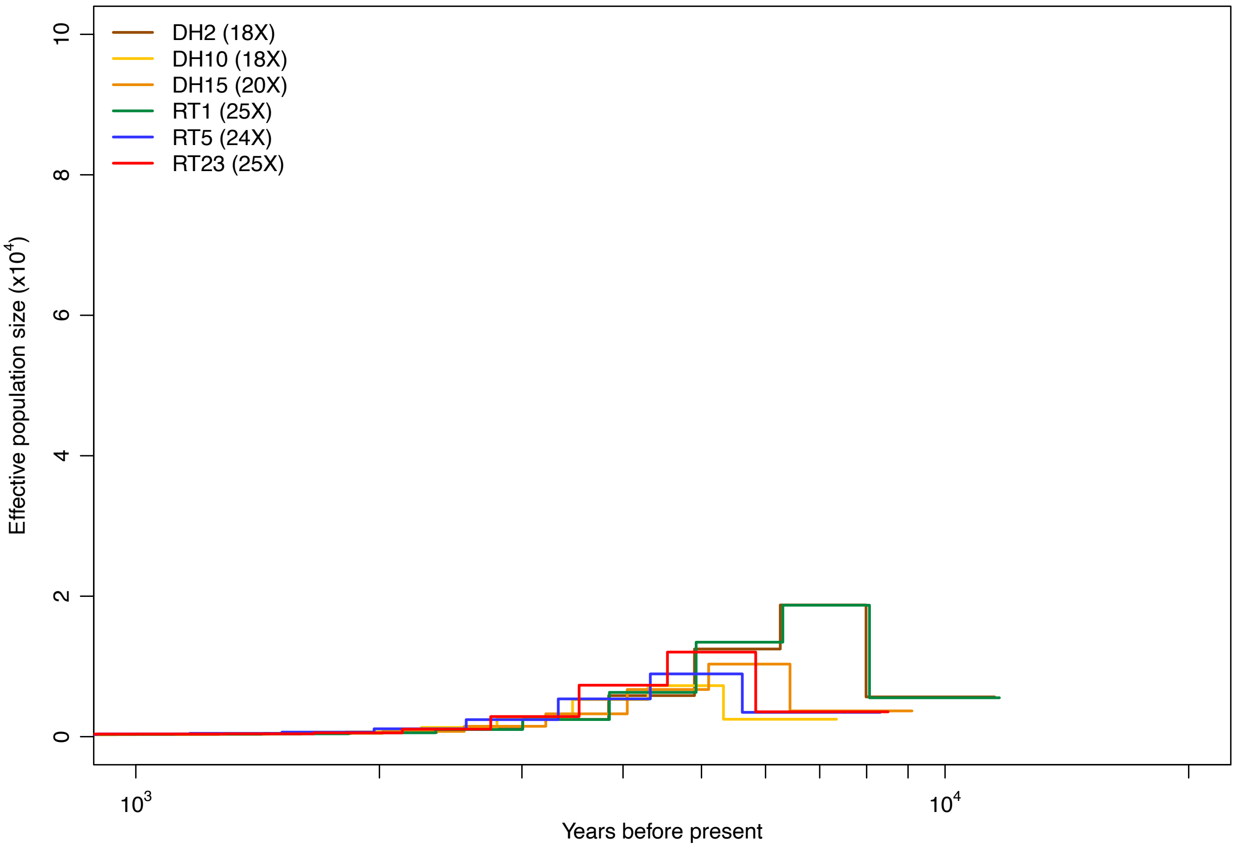


**Supp. Fig. 3**. Zoomed in view of historic effective population size inferred by PSMC for the three highest coverage wild (DH) and captive (RT) individuals. Bootstraps not shown to aid in visibility of each individual trajectory.

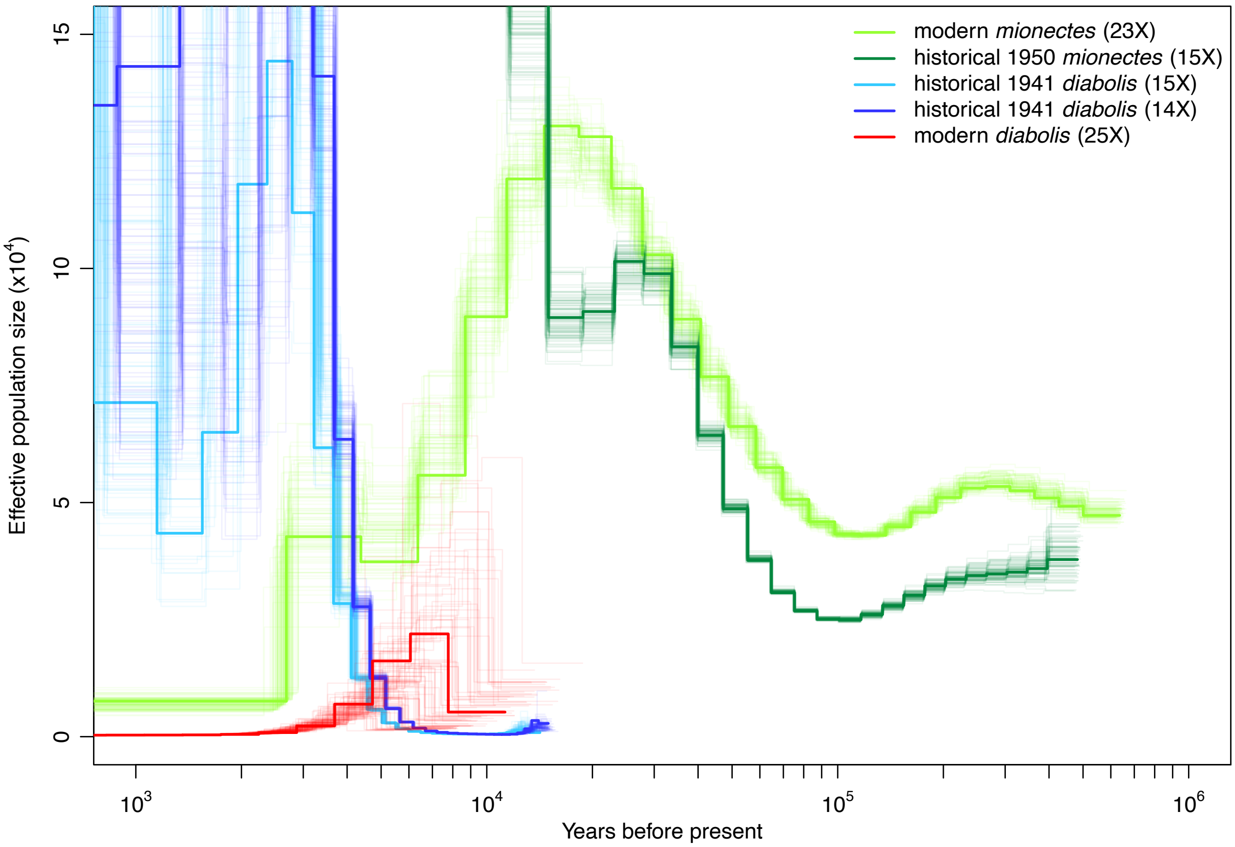


**Supp. Fig. 4.** Comparison of PSMC effective population size trajectories between historical and modern samples of *C. diabolis* and *C. nevadensis mionectes*. Estimates of effective population size on recent timescales are stochastic for the historical genomes likely due to degradation but are not stochastic at deeper timescales. The overall shape of the PSMC trajectories between modern and historical *mionectes* match well on deeper timescales, suggesting that the shape of the PSMC trajectories of the historical *diabolis* samples reflect real signal as well.

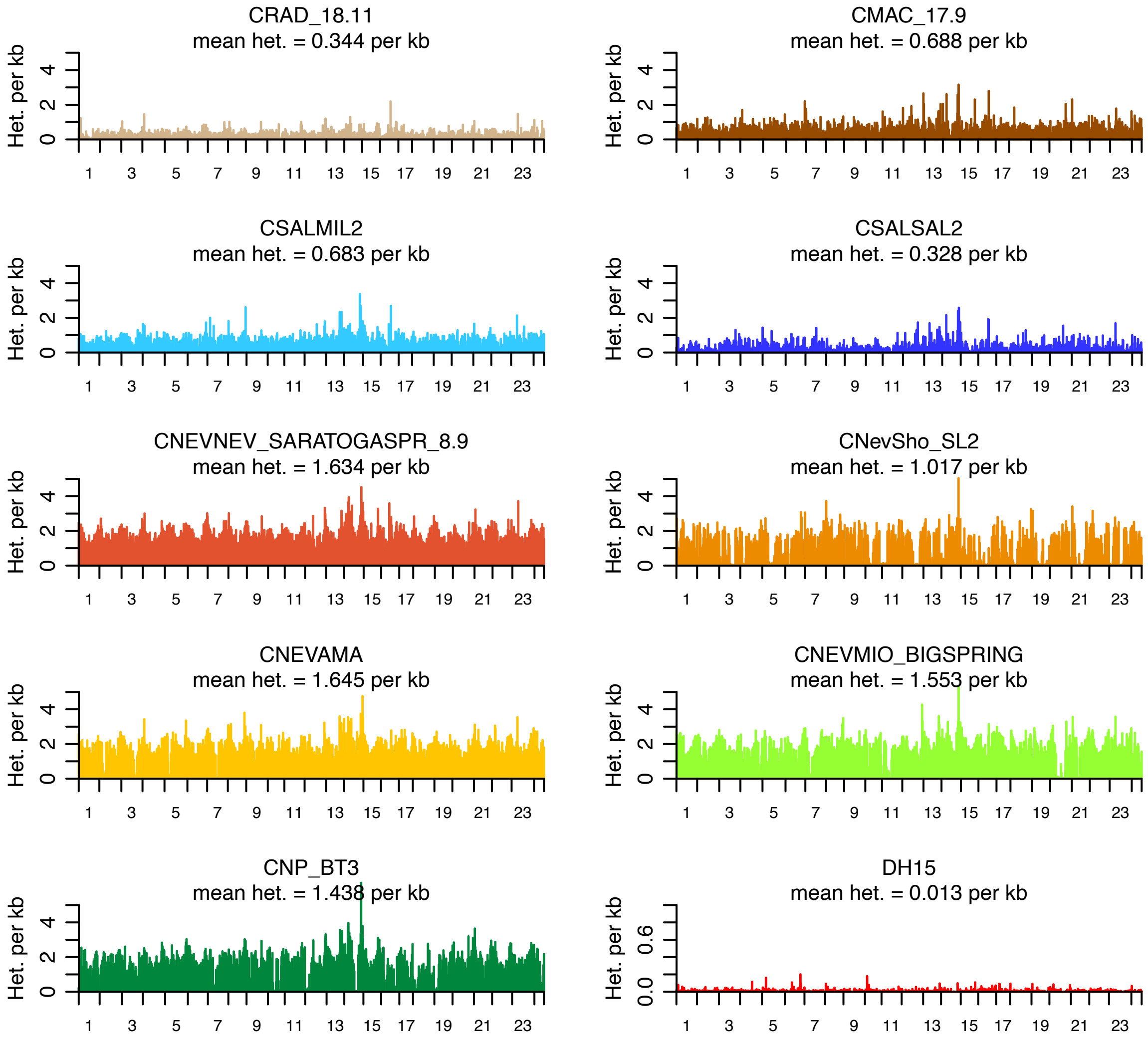


**Supp. Fig. 5.** Bar plots of per-site heterozygosity in non-overlapping 1 Mb windows across the 24 chromosomes for a representative individual per species. We only included windows for which at least half of the genotypes in window were called.

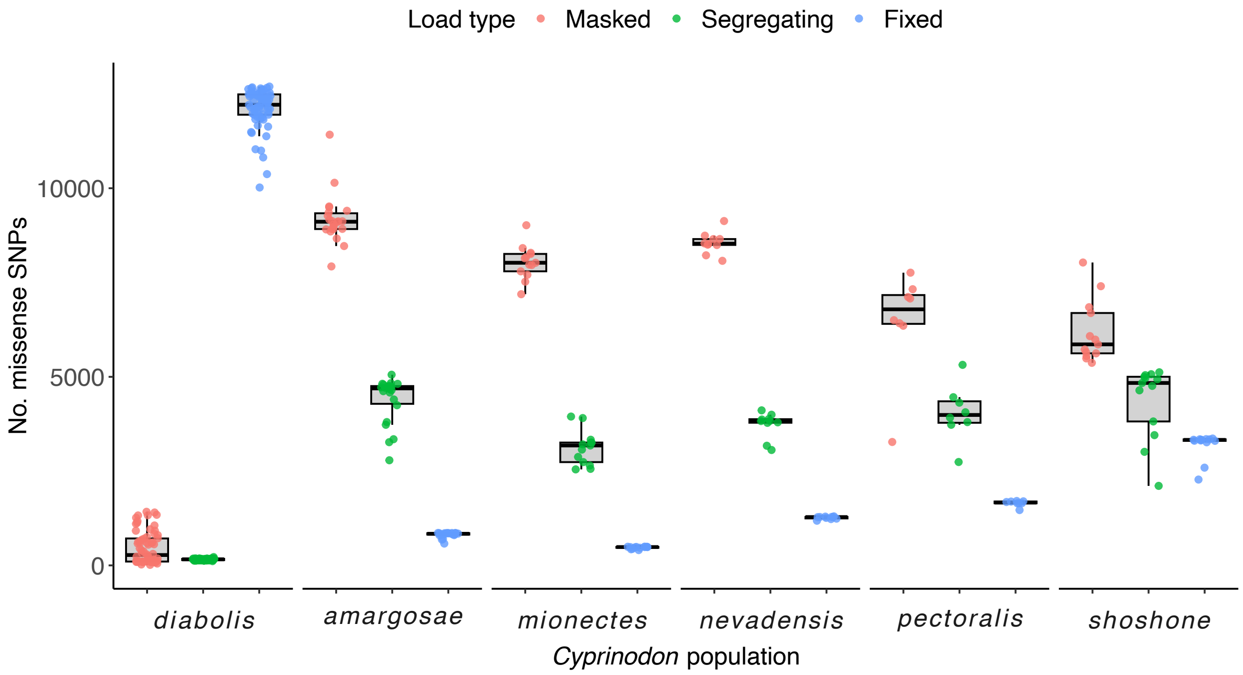


**Supp. Fig. 6.** Individual missense mutation loads based on number of putatively deleterious alleles detected in each individual for Devils Hole pupfish and five *C. nevadensis* populations. These mutations were categorized into the masked load (heterozygous SNPs) and the realized load (homozygous SNPs). The realized load was further parsed into the segregating load (variable within a species) and drift load (fixed within a species or population). Differences in load between species are relative rather than absolute, given that we unable to comprehensively polarize our VCF.

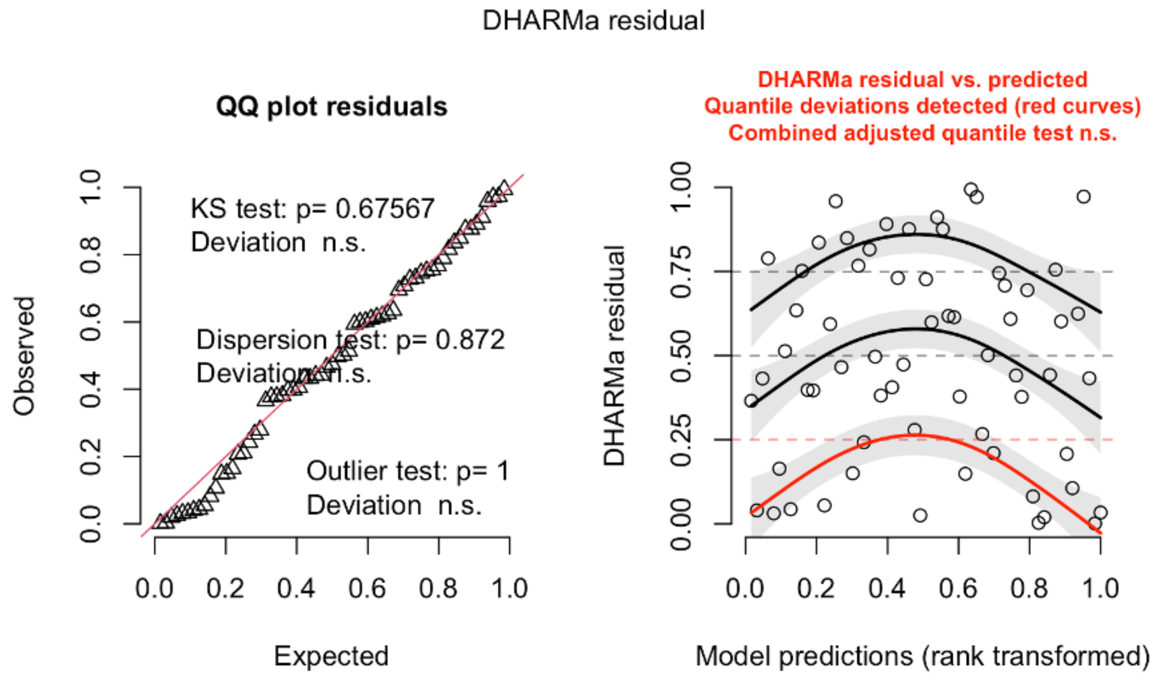


**Supp. Fig. 7.** QQ plot showing uniform distribution of residuals and comparison of residuals vs. model predictions. Residuals show some evidence of systematic deviation, suggesting potential under- or over-prediction in some ranges of values, but not significantly so.

**Supplemental Tables**

**Supp. Table 1.** QUAST output comparing scaffold level assemblies of C_diabolis_v1.0 and C_nevadensis _mionectes_v1.0.

| Assembly | *C. diabolis* reference genome | *C. nevadensis mionectes* reference genome |
|---|---|---|
| # contigs (>= 0 bp) | 1102 | 1726 |
| # contigs (>= 1000 bp) | 1102 | 1726 |
| # contigs (>= 5000 bp) | 987 | 1331 |
| # contigs (>= 10000 bp) | 941 | 1187 |
| # contigs (>= 25000 bp) | 761 | 912 |
| # contigs (>= 50000 bp) | 540 | 684 |
| Total length (>= 0 bp) | 1220346717 | 1216518869 |
| Total length (>= 1000 bp) | 1220346717 | 1216518869 |
| Total length (>= 5000 bp) | 1220112572 | 1215704301 |
| Total length (>= 10000 bp) | 1219783563 | 1214716001 |
| Total length (>= 25000 bp) | 1216731119 | 1210174485 |
| Total length (>= 50000 bp) | 1208951274 | 1202033491 |
| # contigs | 1102 | 1726 |
| Largest contig | 55109245 | 52066102 |
| Total length | 1220346717 | 1216518869 |
| GC (%) | 39.54 | 39.49 |
| N50 (contig) | 48018592 | 46318508 |
| N75 | 42990238 | 39810642 |
| L50 | 13 | 13 |
| L75 | 19 | 20 |
| # N's per 100 kbp | 11.6 | 32.18 |

**Supp. Table 2.** BUSCO (v.5.2.2) scores comparing final assemblies of C_diabolis_v1.0 and C_nevadensis _mionectes_v1.0 using the Augustus gene predictor and the actinopterygii_odb10 lineage database.

| BUSCO Type | *C. diabolis* reference genome | | *C. nevadensis mionectes* reference genome | |
|---|---|---|---|---|
| Complete | 96.1% | 3500 | 97.0% | 3528 |
| Complete and single-copy | 95.2% | 3467 | 95.1% | 3460 |
| Complete and duplicated | 0.9% | 33 | 1.9% | 68 |
| Fragmented | 0.5% | 18 | 0.5% | 18 |
| Missing | 3.4% | 122 | 2.5% | 94 |
| Total BUSCOs searched | 3640 | 3640 | 3640 | 3640 |

**Supp. Table 3.** Effective population sizes based on genome wide linkage-disequilibrium estimated by *currentNe2* (Santiago et al. 2025) for populations sampled at a given time point with at least 5 individuals. Results for *C. sal. salinus* likely did not converge due to low sampling of a relatively large population.

| *Cyprinodon* species | Population | Year | Sample size | Ne estimate (90% CI) |
|---|---|---|---|---|
| *C. diabolis* | Devils Hole | 2019-2020 | 16 | 18.00 (13.67 - 23.70) |
| *C. diabolis* | Ash Meadows Fish Conservation Facility | 2020 | 25 | 18.00 (15.00 - 21.59) |
| *C. macularius* | Coachella Valley | 1994 | 11 | 128.01 (63.63 - 257.53) |
| *C. nev. amargosae* | Tecopa | 1994 | 13 | 25.45 (19.06 - 33.98) |
| *C. nev. amargosae* | main stem of Amargosa River | 2014 | 10 | 4923.4 (505.44 - 47,940.31) |
| *C. nev. mionectes* | Big Spring | 1994 | 6 | 10.40 (7.07 - 15.30) |
| *C. nev. mionectes* | Point of Rocks Spring | 2010 | 7 | 620.68 (137.56 - 2800.48) |
| *C. nev. nevadensis* | Saratoga Spring | 2016 | 6 | 359.64 (90.04 - 1436.46) |
| *C. nev. pectoralis* | School Spring | 2010 | 6 | 16.40 (10.52 - 25.55) |
| *C. nev. shoshone* | Shoshone Spring | 2021 | 8 | 170.86 (72.88 - 400.54) |
| *C. radiosus* | White Mt. Res. Station | 1994 | 11 | 84.4 (48.22 - 147.72) |
| *C. sal. milleri* | Cottonball Marsh | 1994 | 7 | 333.67 (92.93 - 1198.09) |
| *C. sal. salinus* | Salt Creek | 1994 | 8 | Results did not converge |
| *C. sal. salinus* | Salt Creek | 2016 | 10 | Results did not converge |

**Supp. Table 4**. Fixed putative loss-of-function variants unique to *C. diabolis*.

| ID | Gene | Loss-of-function variant type |
|---|---|---|
| CHCHD3 | MICOS complex subunit MIC19 | gained stop codon |
| AVPR1A | Vasopressin V1a receptor | gained stop codon |
| TAOK1 | Serine/threonine-protein kinase TAO1 | gained stop codon |
| col6a6 | Collagen alpha-6(VI) chain | gained stop codon |
| ZYX | Zyxin | gained stop codon |
| PLEC | Plectin | gained stop codon |
| TBC1D7 | TBC1 domain family member 7 | gained stop codon |
| prep | Prolyl endopeptidase | gained stop codon |
| MAP7D1 | MAP7 domain-containing protein 1 | gained stop codon |
| EIF3E | Eukaryotic translation initiation factor 3 subunit E | gained stop codon |
| CCDC126 | Coiled-coil domain-containing protein 126 | gained stop codon |
| C1GALT1 | Glycoprotein-N-acetylgalactosamine 3-beta-galactosyltransferase 1 | gained stop codon |
| RSU1 | Ras suppressor protein 1 | gained stop codon |
| RBM33 | RNA-binding protein 33 | gained stop codon |
| CYP27A1 | Sterol 26-hydroxylase, mitochondrial | gained stop codon |
| L1TD1 | LINE-1 type transposase domain-containing protein 1 | gained stop codon |
| FSTL1 | Follistatin-related protein 1 | gained stop codon |
| DST | Dystonin | gained stop codon |
| RHOQ | Rho-related GTP-binding protein RhoQ | gained stop codon |
| USP54 | Ubiquitin carboxyl-terminal hydrolase 54 | gained stop codon |
| DBN1 | Drebrin | gained stop codon |
| erap2 | Endoplasmic reticulum aminopeptidase 2 | gained stop codon |
| CHST3 | Carbohydrate sulfotransferase 3 | gained stop codon |
| PPP3CC | Serine/threonine-protein phosphatase 2B catalytic subunit gamma isoform | gained stop codon |
| DHX57 | Putative ATP-dependent RNA helicase DHX57 | gained stop codon |
| SCN4A | Sodium channel protein type 4 subunit alpha | gained stop codon |
| ACADSB | Short/branched chain specific acyl-CoA dehydrogenase, mitochondrial | gained stop codon |
| IRF2BPL | Probable E3 ubiquitin-protein ligase IRF2BPL | gained stop codon |
| ACSF2 | Medium-chain acyl-CoA ligase ACSF2, mitochondrial | gained stop codon |
| ADAP2 | Arf-GAP with dual PH domain-containing protein 2 | gained stop codon |
| mchr2 | Melanin-concentrating hormone receptor 2 | lost start codon |
| TRHR | Thyrotropin-releasing hormone receptor | lost start codon |
| CTNND2 | Catenin delta-2 | lost start codon |

| | | |
|---|---|---|
| GRTP1 | Growth hormone-regulated TBC protein 1 | lost start codon |
| MAGI1 | Membrane-associated guanylate kinase, WW and PDZ domain-containing protein 1 | lost stop codon |
| L1TD1 | LINE-1 type transposase domain-containing protein 1 | lost stop codon |
| RAB36 | Ras-related protein Rab-36 | lost stop codon |
| TSTA3 | GDP-L-fucose synthase | lost stop codon |

**Supp. Table 5.** Manually curated deletions identified by DELLY unique to *C. diabolis*.

| ID | Gene | Size | Deletion location | Exon overlaps |
|---|---|---|---|---|
| PFKM | phosphofructokinase | 13835 bp | Chr20: 25725159 - 25738994 | Exons 1-17 |
| RAD23A | UV excision repair protein | 8114 bp | Chr16: 34066402 - 34074516 | Exons 1-3 |

**Supp. Table 6.** Individual sample metadata for newly sequenced individuals in this study.

| Sample ID | Population | Locality | Year |
|---|---|---|---|
| CNevAma_SL1 | *amargosae* | Amargosa River | 2014 |
| CNevAma_SL10 | *amargosae* | Amargosa River | 2014 |
| CNevAma_SL2 | *amargosae* | Amargosa River | 2014 |
| CNevAma_SL3 | *amargosae* | Amargosa River | 2014 |
| CNevAma_SL4 | *amargosae* | Amargosa River | 2014 |
| CNevAma_SL5 | *amargosae* | Amargosa River | 2014 |
| CNevAma_SL6 | *amargosae* | Amargosa River | 2014 |
| CNevAma_SL7 | *amargosae* | Amargosa River | 2014 |
| CNevAma_SL8 | *amargosae* | Amargosa River | 2014 |
| CNevAma_SL9 | *amargosae* | Amargosa River | 2014 |
| CNEVAMA_TECOPA_3-10 | *amargosae* | Tecopa | 1994 |
| CNEVAMA_TECOPA_3-12 | *amargosae* | Tecopa | 1994 |
| CNEVAMA_TECOPA_3-19 | *amargosae* | Tecopa | 1994 |
| CNEVAMA_TECOPA_3-22 | *amargosae* | Tecopa | 1994 |
| CNEVAMA_TECOPA_3-26 | *amargosae* | Tecopa | 1994 |
| CNEVAMA_TECOPA_3-29 | *amargosae* | Tecopa | 1994 |
| CNEVAMA_TECOPA_3-32 | *amargosae* | Tecopa | 1994 |
| CNEVAMA_TECOPA_3-35 | *amargosae* | Tecopa | 1994 |
| CAS_22994-1 | *diabolis* | Devils Hole | 1948 |
| CAS_22994-2 | *diabolis* | Devils Hole | 1948 |
| CAS_22994-3 | *diabolis* | Devils Hole | 1948 |
| CAS_22994-4 | *diabolis* | Devils Hole | 1948 |
| DH1 | *diabolis* | Devils Hole | 2019 |
| DH10 | *diabolis* | Devils Hole | 2020 |
| DH11 | *diabolis* | Devils Hole | 2020 |
| DH12 | *diabolis* | Devils Hole | 2020 |
| DH13 | *diabolis* | Devils Hole | 2020 |
| DH14 | *diabolis* | Devils Hole | 2020 |
| DH15 | *diabolis* | Devils Hole | 2020 |
| DH16 | *diabolis* | Devils Hole | 2020 |
| DH17 | *diabolis* | Devils Hole | 2023 |
| DH18 | *diabolis* | Devils Hole | 2023 |
| DH2 | *diabolis* | Devils Hole | 2019 |
| DH3 | *diabolis* | Devils Hole | 2019 |
| DH4 | *diabolis* | Devils Hole | 2020 |
| DH5 | *diabolis* | Devils Hole | 2020 |

| | | | |
|---|---|---|---|
| DH6 | *diabolis* | Devils Hole | 2020 |
| DH7 | *diabolis* | Devils Hole | 2020 |
| DH8 | *diabolis* | Devils Hole | 2020 |
| DH9 | *diabolis* | Devils Hole | 2020 |
| PR-Dtub1 | *diabolis* | Prop Room | 2023 |
| PR-Dtub10 | *diabolis* | Prop Room | 2023 |
| PR-Dtub11 | *diabolis* | Prop Room | 2023 |
| PR-Dtub12 | *diabolis* | Prop Room | 2023 |
| PR-Dtub2 | *diabolis* | Prop Room | 2023 |
| PR-Dtub3 | *diabolis* | Prop Room | 2023 |
| PR-Dtub4 | *diabolis* | Prop Room | 2023 |
| PR-Dtub5 | *diabolis* | Prop Room | 2023 |
| PR-Dtub6 | *diabolis* | Prop Room | 2023 |
| PR-Dtub7 | *diabolis* | Prop Room | 2023 |
| PR-Dtub8 | *diabolis* | Prop Room | 2023 |
| PR-Dtub9 | *diabolis* | Prop Room | 2023 |
| RT1 | *diabolis* | Refuge Tank | 2020 |
| RT10 | *diabolis* | Refuge Tank | 2019 |
| RT11 | *diabolis* | Refuge Tank | 2020 |
| RT12 | *diabolis* | Refuge Tank | 2020 |
| RT13 | *diabolis* | Refuge Tank | 2020 |
| RT14 | *diabolis* | Refuge Tank | 2020 |
| RT15 | *diabolis* | Refuge Tank | 2020 |
| RT16 | *diabolis* | Refuge Tank | 2020 |
| RT17 | *diabolis* | Refuge Tank | 2020 |
| RT18 | *diabolis* | Refuge Tank | 2020 |
| RT19 | *diabolis* | Refuge Tank | 2020 |
| RT2 | *diabolis* | Refuge Tank | 2020 |
| RT20 | *diabolis* | Refuge Tank | 2020 |
| RT21 | *diabolis* | Refuge Tank | 2020 |
| RT22 | *diabolis* | Refuge Tank | 2020 |
| RT23 | *diabolis* | Refuge Tank | 2020 |
| RT24 | *diabolis* | Refuge Tank | 2020 |
| RT25 | *diabolis* | Refuge Tank | 2020 |
| RT26 | *diabolis* | Refuge Tank | 2023 |
| RT27 | *diabolis* | Refuge Tank | 2023 |
| RT29 | *diabolis* | Refuge Tank | 2023 |
| RT3 | *diabolis* | Refuge Tank | 2023 |
| RT30 | *diabolis* | Refuge Tank | 2023 |

| | | | |
|---|---|---|---|
| RT31 | *diabolis* | Refuge Tank | 2023 |
| RT32 | *diabolis* | Refuge Tank | 2023 |
| RT33 | *diabolis* | Refuge Tank | 2023 |
| RT34 | *diabolis* | Refuge Tank | 2023 |
| RT35 | *diabolis* | Refuge Tank | 2023 |
| RT36 | *diabolis* | Refuge Tank | 2023 |
| RT37 | *diabolis* | Refuge Tank | 2023 |
| RT38 | *diabolis* | Refuge Tank | 2023 |
| RT39 | *diabolis* | Refuge Tank | 2023 |
| RT4 | *diabolis* | Refuge Tank | 2023 |
| RT40 | *diabolis* | Refuge Tank | 2023 |
| RT5 | *diabolis* | Refuge Tank | 2023 |
| RT6 | *diabolis* | Refuge Tank | 2023 |
| RT7 | *diabolis* | Refuge Tank | 2023 |
| RT8 | *diabolis* | Refuge Tank | 2023 |
| RT9 | *diabolis* | Refuge Tank | 2023 |
| UMMZ_134803-15 | *diabolis* | Devils Hole | 1941 |
| UMMZ_134803-17 | *diabolis* | Devils Hole | 1941 |
| CAS_22994-5 | *diabolis* | Devils Hole | 1948 |
| TUL_75085 | *diabolis* | Devils Hole | 1970 |
| UMMZ_134803-11 | *diabolis* | Devils Hole | 1941 |
| CMAC_17-1 | *macularius* | Cochella Valley | 1994 |
| CMAC_17-12 | *macularius* | Cochella Valley | 1994 |
| CMAC_17-13 | *macularius* | Cochella Valley | 1994 |
| CMAC_17-14 | *macularius* | Cochella Valley | 1994 |
| CMAC_17-15 | *macularius* | Cochella Valley | 1994 |
| CMAC_17-16 | *macularius* | Cochella Valley | 1994 |
| CMAC_17-2 | *macularius* | Cochella Valley | 1994 |
| CMAC_17-3 | *macularius* | Cochella Valley | 1994 |
| CMAC_17-6 | *macularius* | Cochella Valley | 1994 |
| CMAC_17-9 | *macularius* | Cochella Valley | 1994 |
| CSALMIL_9-3 | *milleri* | Cottonball Marsh | 1994 |
| CSALMIL_9-4 | *milleri* | Cottonball Marsh | 1994 |
| CSALMIL_9-5 | *milleri* | Cottonball Marsh | 1994 |
| CSALMIL_9-6 | *milleri* | Cottonball Marsh | 1994 |
| CSALMIL_9-7 | *milleri* | Cottonball Marsh | 1994 |
| CSALMIL_9-8 | *milleri* | Cottonball Marsh | 1994 |
| CAS_82806-1 | *mionectes* | Longstreet Spring | 1950 |
| CAS_82806-2 | *mionectes* | Longstreet Spring | 1950 |

| | | | |
|---|---|---|---|
| CNEVMIO_BIGSPR_10-1 | *mionectes* | Big Spring | 1994 |
| CNEVMIO_BIGSPR_10-2 | *mionectes* | Big Spring | 1994 |
| CNEVMIO_BIGSPR_10-6 | *mionectes* | Big Spring | 1994 |
| CNEVMIO_BIGSPR_10-7 | *mionectes* | Big Spring | 1994 |
| CNEVMIO_BIGSPR_11-9 | *mionectes* | Big Spring | 1994 |
| CNM_BT1 | *mionectes* | Point of Rocks | 2010 |
| CNM_BT2 | *mionectes* | Point of Rocks | 2010 |
| CNM_BT3 | *mionectes* | Point of Rocks | 2010 |
| CNM_BT4 | *mionectes* | Point of Rocks | 2010 |
| CNM_BT5 | *mionectes* | Point of Rocks | 2010 |
| CNM_BT6 | *mionectes* | Point of Rocks | 2010 |
| CNM_BT7 | *mionectes* | Point of Rocks | 2010 |
| UMMZ_140460-1 | *mionectes* | Big Spring | 1942 |
| CNEVNEV_SARATOGASPR_8-7 | *nevadensis* | Saratoga Spring | 1994 |
| CNEVNEV_SARATOGASPR_8-8 | *nevadensis* | Saratoga Spring | 1994 |
| CNEVNEV_SARATOGASPR_8-9 | *nevadensis* | Saratoga Spring | 1994 |
| CNevNev_SL1 | *nevadensis* | Saratoga Spring | 2016 |
| CNevNev_SL2 | *nevadensis* | Saratoga Spring | 2016 |
| CNevNev_SL3 | *nevadensis* | Saratoga Spring | 2016 |
| CNevNev_SL4 | *nevadensis* | Saratoga Spring | 2016 |
| CNevNev_SL5 | *nevadensis* | Saratoga Spring | 2016 |
| CNevNev_SL6 | *nevadensis* | Saratoga Spring | 2016 |
| CNEVPEC_SCHOOLSPR_12-14 | *pectoralis* | Warm/School Spring | 1994 |
| CNP_BT1 | *pectoralis* | Warm/School Spring | 1994 |
| CNP_BT2 | *pectoralis* | Warm/School Spring | 1994 |
| CNP_BT3 | *pectoralis* | Warm/School Spring | 1994 |
| CNP_BT4 | *pectoralis* | Warm/School Spring | 1994 |
| CNP_BT5 | *pectoralis* | Warm/School Spring | 1994 |
| CNP_BT6 | *pectoralis* | Warm/School Spring | 1994 |
| C_radiosus_W368-2 | *radiosus* | Well 368 | 2010 |
| C_radiosus_W368-8 | *radiosus* | Well 368 | 2010 |
| CRAD_18-10 | *radiosus* | White Mt. Res. Station | 1994 |
| CRAD_18-11 | *radiosus* | White Mt. Res. Station | 1994 |
| CRAD_18-2 | *radiosus* | White Mt. Res. Station | 1994 |
| CRAD_18-3 | *radiosus* | White Mt. Res. Station | 1994 |
| CRAD_18-4 | *radiosus* | White Mt. Res. Station | 1994 |
| CRAD_18-5 | *radiosus* | White Mt. Res. Station | 1994 |
| CRAD_18-6 | *radiosus* | White Mt. Res. Station | 1994 |
| CRAD_18-7 | *radiosus* | White Mt. Res. Station | 1994 |

| | | | |
|---|---|---|---|
| CRAD_18-8 | *radiosus* | White Mt. Res. Station | 1994 |
| CRAD_18-9 | *radiosus* | White Mt. Res. Station | 1994 |
| CSALSAL_15-10 | *salinus* | Salt Creek | 1994 |
| CSALSAL_15-11 | *salinus* | Salt Creek | 1994 |
| CSALSAL_15-12 | *salinus* | Salt Creek | 1994 |
| CSALSAL_15-15 | *salinus* | Salt Creek | 1994 |
| CSALSAL_15-4 | *salinus* | Salt Creek | 1994 |
| CSALSAL_15-5 | *salinus* | Salt Creek | 1994 |
| CSALSAL_15-7 | *salinus* | Salt Creek | 1994 |
| CSALSAL_15-9 | *salinus* | Salt Creek | 1994 |
| CSalSal_SL1 | *salinus* | Salt Creek | 2016 |
| CSalSal_SL10 | *salinus* | Salt Creek | 2016 |
| CSalSal_SL2 | *salinus* | Salt Creek | 2016 |
| CSalSal_SL3 | *salinus* | Salt Creek | 2016 |
| CSalSal_SL4 | *salinus* | Salt Creek | 2016 |
| CSalSal_SL5 | *salinus* | Salt Creek | 2016 |
| CSalSal_SL6 | *salinus* | Salt Creek | 2016 |
| CSalSal_SL7 | *salinus* | Salt Creek | 2016 |
| CSalSal_SL8 | *salinus* | Salt Creek | 2016 |
| CSalSal_SL9 | *salinus* | Salt Creek | 2016 |
| LACM_25258-1 | *salinus* | Salt Creek | 1978 |
| LACM_25264-1 | *salinus* | Salt Creek | 1981 |
| CNEVSHO_HEADSPR_6-11 | *shoshone* | Shoshone, Head Springs Pool | 1994 |
| CNEVSHO_HEADSPR_6-3 | *shoshone* | Shoshone, Head Springs Pool | 1994 |
| CNEVSHO_HEADSPR_6-4 | *shoshone* | Shoshone, Head Springs Pool | 1994 |
| CNevSho_SL1 | *shoshone* | Shoshone Spring | 2021 |
| CNevSho_SL2 | *shoshone* | Shoshone Spring | 2021 |
| CNevSho_SL3 | *shoshone* | Shoshone Spring | 2021 |
| CNevSho_SL4 | *shoshone* | Shoshone Spring | 2021 |
| CNevSho_SL5 | *shoshone* | Shoshone Spring | 2021 |
| CNevSho_SL6 | *shoshone* | Shoshone Spring | 2021 |
| CNevSho_SL7 | *shoshone* | Shoshone Spring | 2021 |
| CNevSho_SL8 | *shoshone* | Shoshone Spring | 2021 |
